# The aminoalkylindole, BML-190, negatively regulates chitosan synthesis via the cAMP/PKA1 pathway in *Cryptococcus neoformans*

**DOI:** 10.1101/751172

**Authors:** Brian T. Maybruck, Woei C. Lam, Charles A. Specht, Ma Xenia G. Ilagan, Maureen J. Donlin, Jennifer K. Lodge

## Abstract

*Cryptococcus neoformans* can cause fatal meningoencephalitis in patients with AIDS or other immune-compromising conditions. Current antifungals are suboptimal to treat this disease, therefore, novel targets and new therapies are needed. Previously, we have shown that chitosan is a critical component of the cryptococcal cell wall, is required for survival in the mammalian host, and that chitosan deficiency results in rapid clearance from the mammalian host. We had also identified several specific proteins that were required for chitosan biosynthesis, and we hypothesize that screening for compounds that inhibit chitosan biosynthesis would identify additional genes/proteins that influence chitosan biosynthesis.

To identify these compounds we developed a robust and novel cell-based flow cytometry screening method to identify small molecule inhibitors of chitosan production. We screened the ICCB Known Bioactives library and identified 8 compounds that reduced chitosan in *C. neoformans*. We used flow cytometry-based counter and confirmatory screens, followed by a biochemical secondary screen to refine our primary screening hits to 2 confirmed hits.

One of the confirmed hits that reduced chitosan content was the aminoalkylindole, BML-190, a known inverse agonist of mammalian cannabinoid receptors. We demonstrated that BML-190 likely targets the C. neoformans G-protein coupled receptor, Gpr4, and via the cAMP/PKA signaling pathway, contributes to an intracellular accumulation of cAMP that results in decreased chitosan. Our discovery suggests that this approach could be used to identify additional compounds and pathways that reduce chitosan biosynthesis, and could lead to potential novel therapeutics against C. neoformans.

**Importance:** *Cryptococcus neoformans* is a fungal pathogen that kills ∼200,000 people every year. The cell wall is an essential organelle that protects fungus from the environment. Chitosan, the deacetylated form of chitin, has been shown to be an essential component of cryptococcal cells wall during infection of a mammalian host. In this study, we screened a set of 480 compounds, which are known to have defined biological activities, for activity that reduced chitosan production in *C. neoformans*. Two of these compounds were validated using an alternative method of measuring chitosan, and one of these was demonstrated to impact the cAMP signal transduction pathway. This work demonstrates that the cAMP pathway regulates chitosan in *C. neoformans*, and validates that this screening approach could be used to find potential antifungal agents.

## Introduction

Despite the current antifungal therapies, up to 19% of AIDS-related deaths (∼200,000) occur globally due to cryptococcal meningitis (1–3). Several factors contribute to this high mortality rate including i) toxicities from current therapies such that many patients cannot tolerate the regime (2–3), ii) relative expense and difficulty of administering some of the formulations, and iii) the long duration of the regime reduces patient compliance. Although the implementation of antiretroviral therapy (ART) has been profound in its success of reducing AIDS-related deaths (1), to avoid immune reconstitution inflammatory syndrome (IRIS) that is linked to ART, ART must be delayed until successful antifungal therapy can be first applied (4–5). With the suboptimal efficacy of current antifungal therapies it can be challenging to avoid the effects of IRIS. Furthermore, antimicrobial drug resistance is an identified concern when using the current antifungal therapies against cryptococcosis (6–7). Overall, these concerns create the urgency for the discovery of novel targets for the development of effective drugs against the primary etiology of AIDS-related cryptococcal meningitis, *Cryptococcus neoformans*.

Chitosan is a key component of the *C. neoformans* cell wall and is essential for fungal survival in the mammalian host (8–9). The precursor to chitosan in *C. neoformans* is chitin. Chitin is synthesized by one or more of the integral membrane proteins, chitin synthases (Chs), some of which require an accessory protein (Csr), from cytoplasmic stores of uridine diphosphate N-acetylglucosamine and is then deacetylated by chitin deacetylases (Cda) to produce chitosan. Although most fungi have genes encoding putative chitin deacetylases, chitosan has been shown to be present in vegetatively growing cells of a small subset of fungi, and chitosan biosynthesis has only been studied in a few species. Chitin is made up of beta-(1, 4)-linked *N*-acetylglucosamine (GlcNAc) monomers, and when greater than 50% of these monomers are deacetylated by Cda’s to glucosamine (GlcN; 8-9), the polymer becomes chitosan. Cryptococcus has eight chitin synthase genes, and only one of them (*CHS3*) affects chitosan production (10). Deletion of the *CSR2* gene which encodes the accessory protein suggested to be associated with Chs3 also prevents chitosan production. The chitin produced by Chs3/Csr2 is subsequently deacetylated into chitosan by three chitin deacetylases: Cda1, Cda2, and Cda3 (9–10).

Three *C. neoformans* strains have been engineered to be deficient in chitosan production, via gene deletions of *CHS3*, *CSR2*, or by deletion of all three *CDA* genes. Chitosan in these three strains was undetectable or very low by biochemical assay or using a chitosan specific stain (9–10). The phenotypes of these strains demonstrated that chitosan is important for: i) cell wall integrity, ii) cell budding, iii) cell size and morphology, and iv) virulence (8–9). Previously we demonstrated that in a mouse inhalation model of infection, chitosan-deficient strains are avirulent and can be cleared within two days (8). We examined the host immune response to infection with the *cda1*Δ*cda2*Δ*cda3*Δ chitosan-deficient strain and found an increase in the Th1-mediated immune response. Mice infected with this strain were also fully protected from subsequent challenge with wild-type KN99 (11). We, along with other groups, have shown that these immunological responses are necessary for protection from subsequent *C. neoformans* infections (12–13). Taken together, these results identify chitosan as being important for *C. neoformans* virulence.

Based on the rapid clearance of the *cda1*Δ*cda2*Δ*cda3*Δ chitosan-deficient strain, we hypothesize that compounds that interfere with chitosan biosynthesis and cause chitosan deficiency could facilitate clearance of *C. neoformans* and these compounds would provide clues to additional pathways and proteins that regulate or contribute to chitosan production. Therefore, we developed a novel, medium-throughput, cell-based, phenotypic flow cytometry screening assay to identify small molecules that reduce chitosan levels. We screened the Institute of Chemistry and Cell Biology (ICCB) known bioactives library and discovered that the aminoalkylindole, BML-190, targets the *C. neoformans* G-protein coupled receptor, Gpr4, and via the cAMP/PKA signaling pathway, contributes to an intracellular accumulation of cAMP that results in decreased chitosan.

## Results

### A novel medium-throughput flow cytometry assay that robustly measures chitosan levels by BR fluorescence

To identify small molecules that can inhibit chitosan production, we developed a cell-based phenotypic medium-throughput flow cytometry screen that is sensitive and robust (Z-prime >0.6) in detecting *C. neoformans* chitosan levels using the chitosan selective fluorescent dye Cibacron Brilliant Red (BR; Fig. S1A; 18-19).

We confirmed that this flow cytometry-based assay reliably detects differences in chitosan content of strains of Cryptococcus by comparing it to the low-throughput 3-methyl-2-benzothiazolone hydrazone hydro-chloride (MBTH) chitosan assay (20). We found a statistically significant positive correlation between the fluorescence from BR stained *C. neoformans* strains that differentially produce chitosan, measured via flow cytometry compared to the levels of chitosan in those same strains, as measured by the MBTH assay (Fig. S1B-C). The key advantages of the flow cytometry-based assay over the MBTH assay are its miniaturized, microplate format (96-well), increased throughput and the ability to measure multiple parameters from a single sample/well.

### Screening strategy to identify small-molecule inhibitors of chitosan synthesis in *C. neoformans*

We chose to screen the 480 compound Institute of Chemistry and Cell Biology (ICCB) Known Bioactives library because the biological targets of the compounds in this library are known, allowing us to potentially identify the mechanism of action of our hits. We developed a screening strategy funnel that uses a series of assays that are increasingly sensitive in their ability to identify small-molecule inhibitors of chitosan production so primary hits from the flow cytometry assay could be confirmed by orthogonal approaches (Fig. S2).

There are five proteins that are already known to affect chitosan production – Chs3, Csr2, Cda1, Cda2 and Cda3 (8–10). Ideally, the screen would identify inhibitors of each of these proteins, along with other, as yet unknown, proteins or pathways. However, the Cdas are functionally redundant, making it less likely that the screen would identify a deacetylase inhibitor. Therefore, we did our initial screening with a strain that expressed only the *CDA1* gene. We chose *CDA1* because it is upregulated in the mouse upon infection and deletion of *CDA1* results in avirulence, while deletion of both *CDA2* and *CDA3* maintains virulence similar to wild-type (21). Therefore, for screening, we chose a strain that expresses *CDA1*, but has both *CDA2* and *CDA3* deleted (i.e., *cda2Δ3Δ*). The *cda2Δ3Δ* strain produces sufficient chitosan (Fig. S1), retains a wild-type morphology for our screens, and has all the remaining elements of the chitosan production machinery intact, so that we should still be able to identify inhibitors of other factors that impact chitosan production from our screens. The negative control for this screen was untreated *cda2Δ3Δ* strain and the positive control was the chitosan-deficient *cda1Δ2Δ3Δ* strain.

To minimize concerns associated with the viability of screening strains, prior to subjecting them to the primary screening conditions, they were first grown in the optimal growth liquid medium yeast extract-peptone-dextrose (YPD) for 3 days at 30°C under constant shaking (300 RPM). Next, they were washed 2x in PBS and then added at 5 × 10^5^ cells/ml to 96-well round bottom plates containing 100 μl cRPMI (0.625% HI-FBS) and 1 μl of the ICCB library compound for a 1% DMSO final concentration with varied compound concentrations (i.e., 1μM, 10μM, 5μg/ml, or 50 μg/ml). The varied compound concentrations were based upon what was provided by the supplier. Furthermore, the ICCB library compounds were screened in duplicate to help reduce the likelihood of false-negatives. In the context of growth conditions considered, if compounds do have an effect on altering chitosan levels, for them to even have the potential for pre-clinical to clinical applications, they would ultimately need to do so under host conditions. Therefore, we chose to characterize their effects under conditions that they would most closely simulate that of a mammalian host (i.e., complete RPMI, 5% CO_2_, and 37°C). After 3 days under tissue culture conditions, cells were stained with BR and then fixed in 1% paraformaldehyde (PFA) (Fig 1A).

**Figure 1.**
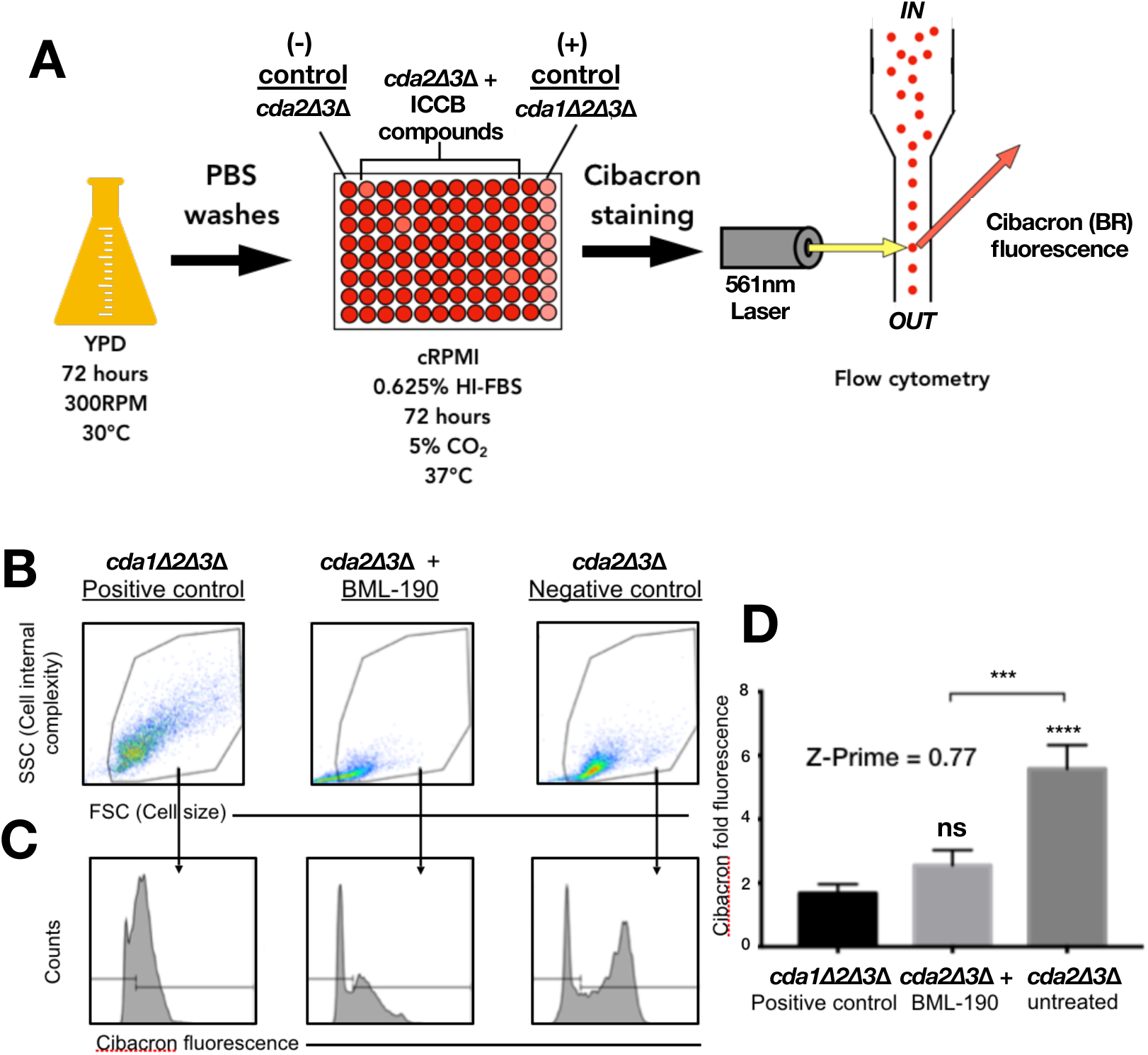
Primary screening strategy. A) An overview of the primary cell-based phenotypic screen of the 480 compounds from the ICCB library. B) From each well, 7500 cell events were collected from gated region. C) The gated region was further analyzed for gMFI BR fluorescence. D) Upon comparison to controls, BML-190, was identified as a tentative ‘hit’. Cibacron (BR) fold fluorescence = geometric mean fluorescence intensity (gMFI) of BR stained, drug + cda2Δ3Δ/gMFI of unstained *cda2Δ3Δ* strain. Means were compared to the positive control using a one-way ANOVA followed by the Bonferroni multiple-comparison test. A t-test was used to show a difference between our screening strain with/without drug. ***, p<0.001; and ***, p<0.0001.

Plates were then analyzed for multiple parameters by flow cytometry. Specifically, from each well, 7500 cell events were collected from a gated region, within the forward (FSC) vs side scatter (SSC) dot plots that predicted morphological phenotypes of *C. neoformans* cells associated with chitosan-deficiency (using our chitosan-deficient strain *cda1Δ2Δ3Δ*, as a guide) and excluded smaller cells that we have empirically determined do not inherently fluoresce BR (Fig 1B). Ultimately, we simultaneously assessed three parameters to aggressively reduce false-positive hits: Cell size (i.e., FSC; proportional to cell-surface area and is diffracted incident light around the cell detected in the forward direction by a photodiode), cellular complexity (i.e., SSC; proportional to the biomolecule complexity of a cell that is refracted incident light that is collected at 90° and then sent to a photodiode), and BR fluorescence. The first two parameters correspond to the initial gated region described above (Fig 1B). The third parameter was assessed from the gated population to measure BR fluorescence (Fig 1C). The assessment of the amount of BR fluorescence from the screening strain within each well was calculated by determining the BR geometric mean fluorescence intensity (gMFI) from the samples. The gMFI was then used to calculate BR fold fluorescence by dividing those numbers by the gMFI of our unstained *cda2Δ3Δ* sample on each plate to quantify the relative levels of chitosan production upon exposure of *C. neoformans* strains to different compounds (Fig 1D). This medium throughput screen (12 plates) was completed in ∼ 6-hours, and had an average plate Z-prime of 0.77, providing us confidence that we have an efficient and robust assay.

We identified initial hits based upon the following criteria: The compound must induce the *cda2Δ3Δ* strain to have a BR fold fluorescence that is: i) statistically different and ii) at least ∼2-fold lower in signal than the untreated *cda2Δ3Δ* screening strain (Fig 1D). This resulted in the identification of eight hits from the ICCB library (Table 1). Based upon the stringency of our hit criteria, this equated to a hit rate of 1.7% which is below what is commonly seen for the screening of the ICCB library (i.e., 2% to 15%; 22-26).

**Table 1.** Primary screen ICCB Bioactives library hits.

| Name | Classification | Known Mode of Action |
| --- | --- | --- |
| BML-190 | Bioactive lipids | Cannabinoid CB1/CB2 inverse agonist |
| DL-PDMP | Lipid biosynthesis | Glucosylceramide synthase inhibitor |
| Quercetin | Kinase inhibitors | Kinase inhibitor |
| Dipyridamole | Inhibitors | cGMP phosphodiesterase inhibitor |
| HA1077 | Kinase inhibitors | Inhibitor of Rho-dependent kinases |
| Juglone | Inhibitors | PIN1 inhibitor |
| Grayanotoxin III | Ion channel ligands | Sodium channels |
| RK-682 | Inhibitors | VHR phosphatase inhibitor |

One effect that the initial hit compounds could have on screening strains is causing their relative reduced growth or cell death of the *cda2Δ3Δ* strain, unrelated to chitosan deficiency. We anticipated that compounds that recapitulate the phenotype of our chitosan-deficient *cda1Δ2Δ3Δ* strain might have similar, modest growth defects, but should grow as well or better than the *cda1Δ2Δ3Δ* strain. Compounds that inhibited the growth more than the *cda1Δ2Δ3Δ* strain are likely acting through a different mechanism than chitosan deficiency and were eliminated. We also repeated the assay used for the primary screen to generate a dose-response profile for each hit. However, two of the eight compounds, grayanotoxin III and RK-682 were eliminated from this analysis due to their high costs compared to other six hits; these two compounds will be assessed further in the future. Hits were assessed for growth inhibition by the inclusion of fluorescent microsphere counting beads into wells with the *cda2Δ3Δ* cells and hit ICCB compounds. This assay was used to define a fourth flow cytometry parameter, cell proliferation. Dose response and proliferation validation were done as described above. We validated 2 of the 6 remaining hits (Table 1), BML-190 (Fig 2B-C) and dipyridamole (Fig. S3B-C).

**Figure 2.**
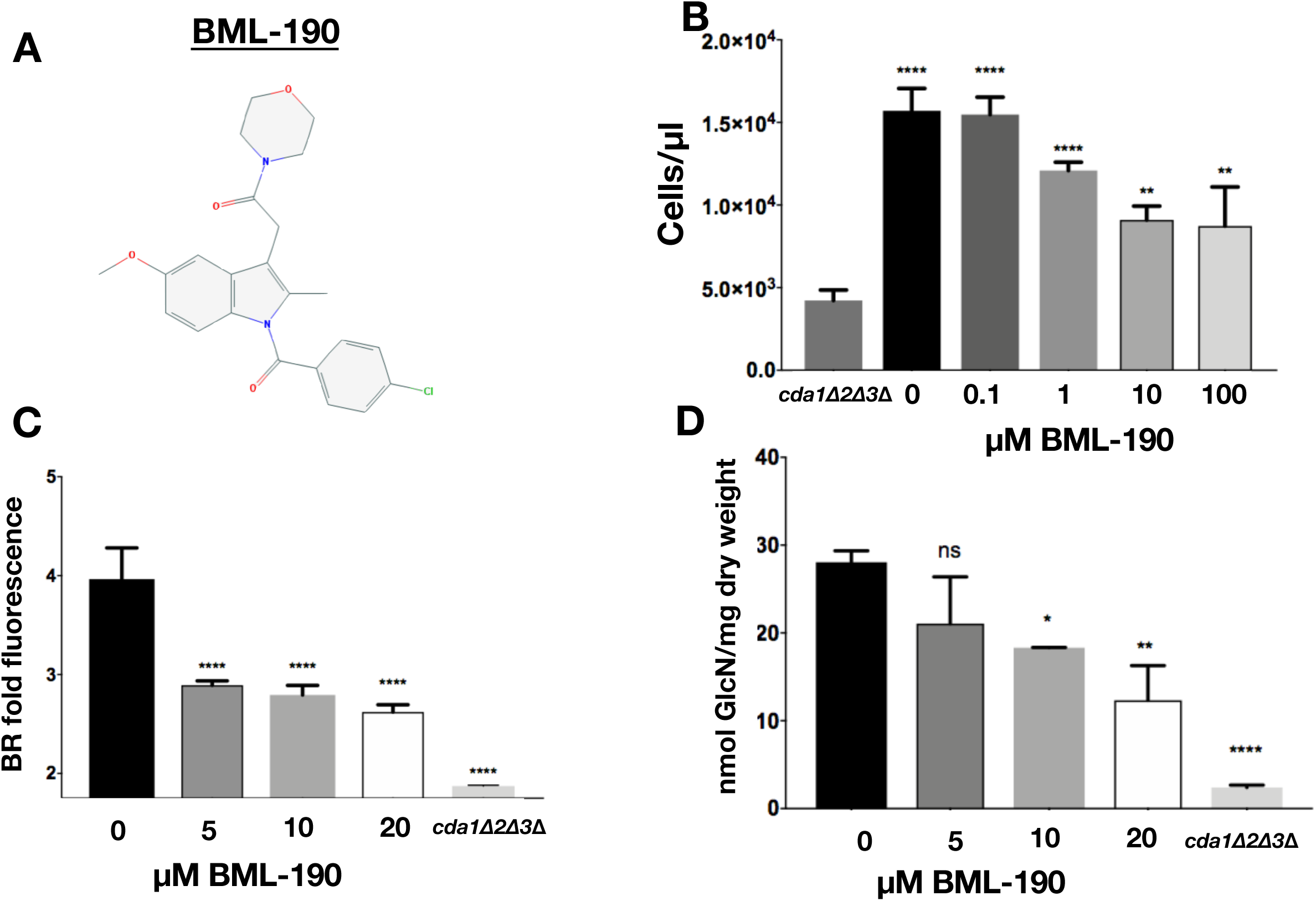
The cannabinoid inverse-agonist, BML-190 decreases chitosan levels. A) Chemical structure of BML-190 (indomethacin morpholinylamide) an aminoalkylindole. B) *C. neoformans* concentrations determined by flow cytometry using counting beads with BML-190 concentrations between 0.1 µM to 100 µM. C) Effect of BML-190 on the chitosan levels of wild-type *C. neoformans* as measured by BR. BR-fold fluorescence = geometric mean fluorescence intensity (gMFI) of BR stained, BML-190 + KN99α/gMFI of unstained KN99α (n = 3). D) Effect of varying doses of BML-190 on the chitosan content as measured by the MBTH assay (n = 3). Means among groups were compared to cda1Δ2Δ3Δ (B) or KN99α (C-D) with no BML-190 added using a one-way ANOVA followed by the Bonferroni multiple-comparison test. *, p<0.05; **, p<0.01; and ****, p<0.0001.

### The aminoalkylindole, BML-190, reduces chitosan synthesis in *C. neoformans*

Our secondary screen used the MBTH biochemical assay to measure chitosan, and both BML-190 and dipyridamole reducing chitosan levels (Fig. 2D and Fig. S3D) using this secondary, biochemical screen.BML-190 is known as an inverse agonist of G-protein coupled receptors (GPCRs) (29). To our knowledge, there was no known connection between GPCRs and chitosan production in *C. neoformans*; thus, we chose to further characterize the effect of BML-190 in *C. neoformans* and to explore the potential role of GPCR signaling in the synthesis of chitosan.

### BML-190 reduces chitosan production in multiple research and clinical strains of *C. neoformans*

*C. neoformans* var. *grubii* (serotype A) strains are important contributors to cryptococcosis in humans (27). Two congenic mating strains from this group, KN99*α* and KN99a, along with multiple clinical strains: TDY1989 and TDY2001 were treated with 10 µM BML-190, and all demonstrated a reduction in their chitosan production (Fig 3). However, as expected, no chitosan was detectable with *Saccharomyces cerevisiae* or our *cda1Δ2Δ3Δ* strain that are both chitosan-deficient. This helps further support the specificity of BR to measuring chitosan since *S. cerevisiae* is known to only produce chitosan during sporulation (28) and we have empirically shown chitosan-deficiency with our *cda1Δ2Δ3Δ* strain.

**Figure 3.**
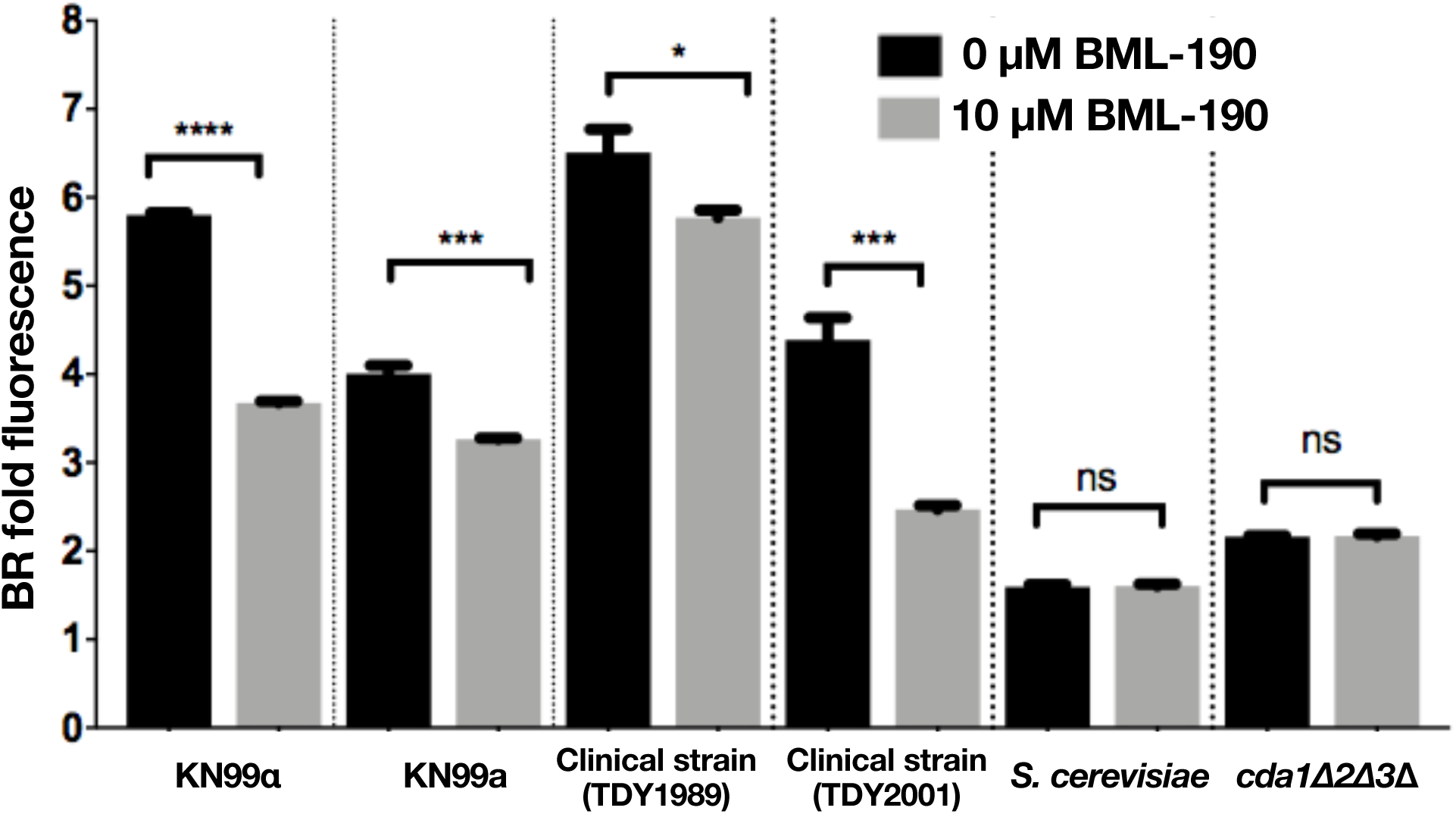
BML-190 causes chitosan reduction in multiple strains. *C. neoformans* strains were cultured with/without BML-190 (10 µM) for 3 days, stained with BR and measured for gMFI by flow cytometry (n = 3). A Student’s t-test was used to show a difference between strains with/without drug. *, p<0.05; ***, p<0.001; and ****, p<0.0001.

### BML-190 acts through the G protein-coupled receptor 4 (Gpr4) of *C. neoformans*

In human cells, BML-190 targets the G protein-coupled receptors (GPRC) cannabinoid receptors 1 and 2 (CNR1/2), with a greater affinity towards CNR2 (29). GPCRs are a family of receptors that are known to be activated by external molecules that are well-known to induce signal transduction in the cAMP pathway, which is important for many cellular responses. BML-190 acts as an inverse agonist of this receptor, resulting in increased cAMP levels. Since G protein signaling is known to be conserved among organisms, with particular importance associated with fungal virulence (30), we examined if BML-190’s mode of action could also target GPCRs from *C. neoformans* (31). Initially, we looked for the closest homologs to CNR1/2 in *C. neoformans* using BLAST and multiple sequence alignments. We identified four, *GPR2*, *GPR3, GPR4* and *GPR5*, although none have more than 30% similarity to the human CNR1/2. To identify the target of BM-190, we used strains that were deleted for each of the four GPCRs has been deleted. These deletion strains were available from the Madhani UCSF *C. neoformans* knockout collection obtained from the Fungal Genetics Stock Center (15). After culturing these strains with BML-190 in cRPMI for 3 days under tissue culture conditions, we discovered that of these four, only the *gpr4Δ* strain showed no decrease in chitosan in response to BML-190 (Fig 4A; Fig. S4). Because the *gpr4Δ* strain was nonresponsive to BML-190, Gpr4 may be the primary receptor for BML-190 in *C. neoformans*.

**Figure 4.**
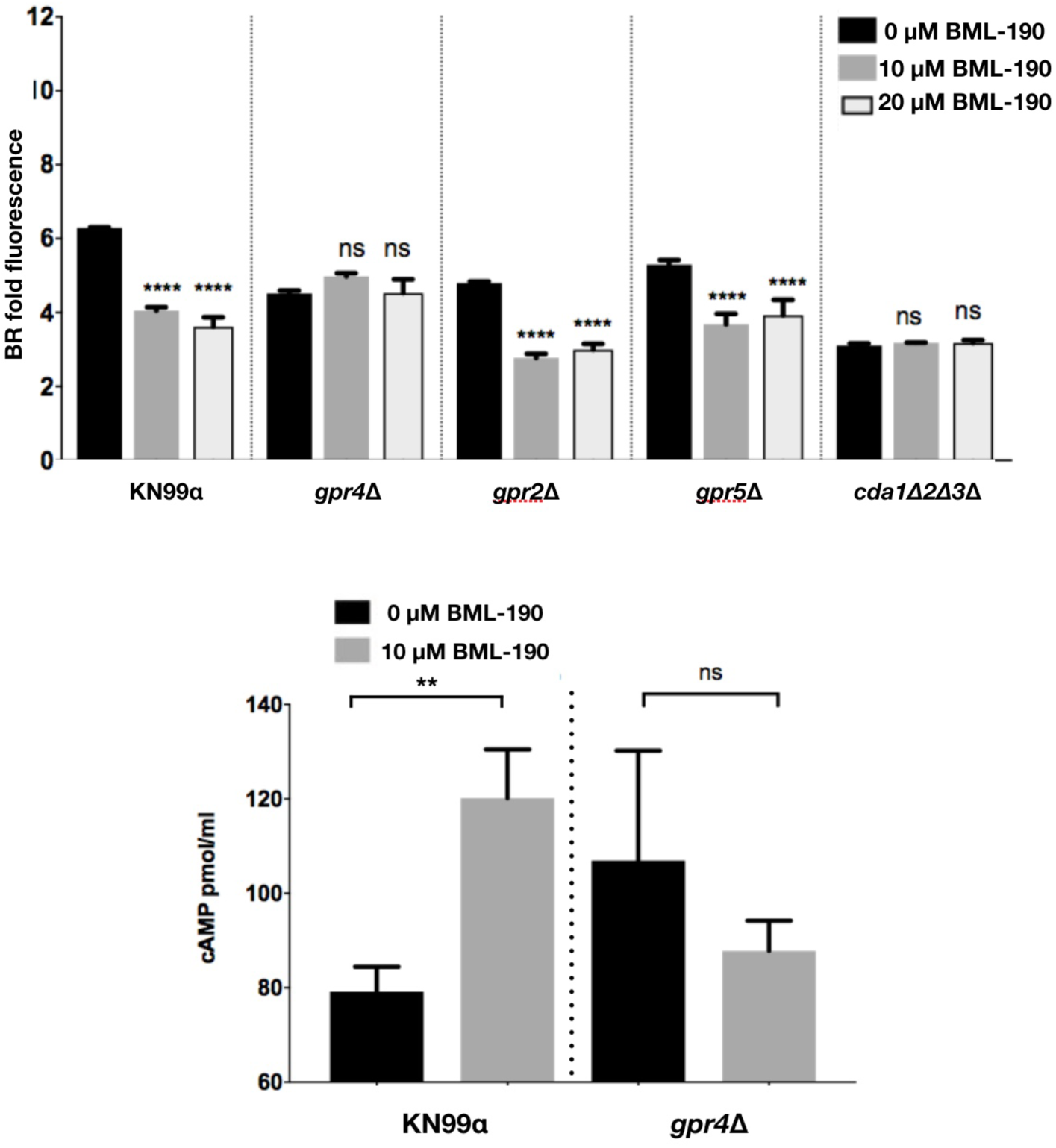
The G-protein coupled receptor Gpr4 is necessary for BML-190 to reduce chitosan levels. A) The *C. neoformans* gpr-deficient strains were cultured with/without BML-190 (10 µM and 50 µM) for 3 days, stained with BR and measured for gMFI by flow cytometry (n = 4). B) BML-190 induces intracellular accumulation of cAMP in wild-type *C. neoformans*, but not in a *gpr4*Δ strain. Means among groups were compared to strains with no BML-190 added using a one-way ANOVA followed by the Bonferroni multiple-comparison test or a Student’s t-test. *, p<0.05; **, p<0.01.****, p<0.0001.

Agonists of cannabinoid receptors are known to contribute to the degradation of intracellular cAMP (33). Similarly, in *C. neoformans*, Gpr4 activates the cAMP/PKA pathway upon stimulation by the agonists glucose and the amino acid methionine that ultimately results in cAMP degradation (30, 34). Inversely, if BML-190 is targeting Gpr4, and acting as an inverse-agonist, then it should cause a build-up of cAMP in wild-type cells, but not in the *gpr4Δ* strain. Therefore, we determined the intracellular content of cAMP in wild-type *C. neoformans* and the *gpr4Δ* strain with and without 10 µM BML-190. We incubated strains with/without BML-190 for 30 minutes, processed cells to stably collect cAMP and demonstrated that BML-190 causes an increase in intracellular cAMP from wild-type cells, but not the *gpr4Δ* strain (Fig 4B). Based on these results, we hypothesize that BML-190 is targeting the Gpr4 and causes an increase in *C. neoformans* intracellular cAMP that inversely correlates with a decrease in chitosan.

### BML-190 reduces chitosan by increasing cAMP levels via the cAMP/PKA pathway

Gpr4 is known to be a receptor in *C. neoformans* for the cAMP/PKA pathway (30, 34). Therefore, we hypothesized that BML-190 is increasing cAMP in wild type Cryptococcus by acting as an inverse agonist to the cAMP/PKA pathway, which is responsible for regulating cAMP (34). To test that idea, we selected three additional strains with deletions of genes in this pathway, including deletions of *GPA1*, *CAC1* and *PKA1*. Gpa1 is the G protein alpha subunit of the heterotrimeric G protein complex that contains Gpr4. Upon ligand activation, Gpr4 activates Gpa1 and dissociates it from the complex to continue the signal transduction cascade binding to the adenylyl cyclase (Cac1), which converts ATP to cAMP. Based upon our hypothesis that cAMP levels are inverse to chitosan, the deletion of either *GPA1* or *CAC1* would be predicted to decrease cAMP production, and thus increase chitosan. Following the incubation of these different strains for 3 days in cRPMI under tissue culture conditions, the *gpa1Δ* and *cac1Δ* strains both showed an increase in chitosan production compared to wild-type and increasing doses of BML-190 had no effect on chitosan in these deletion strains (Fig 5). We also confirmed that addition of BML-190 to the *cac1Δ* strain does not affect chitosan levels, as measured by the MBTH biochemical assay (data not shown). Photomicrographs of BR stained wild-type (KN99*α*) and mutant strains with/without 10 µM BML-190 further confirmed our flow cytometry analysis (Fig 6).

**Figure 5.**
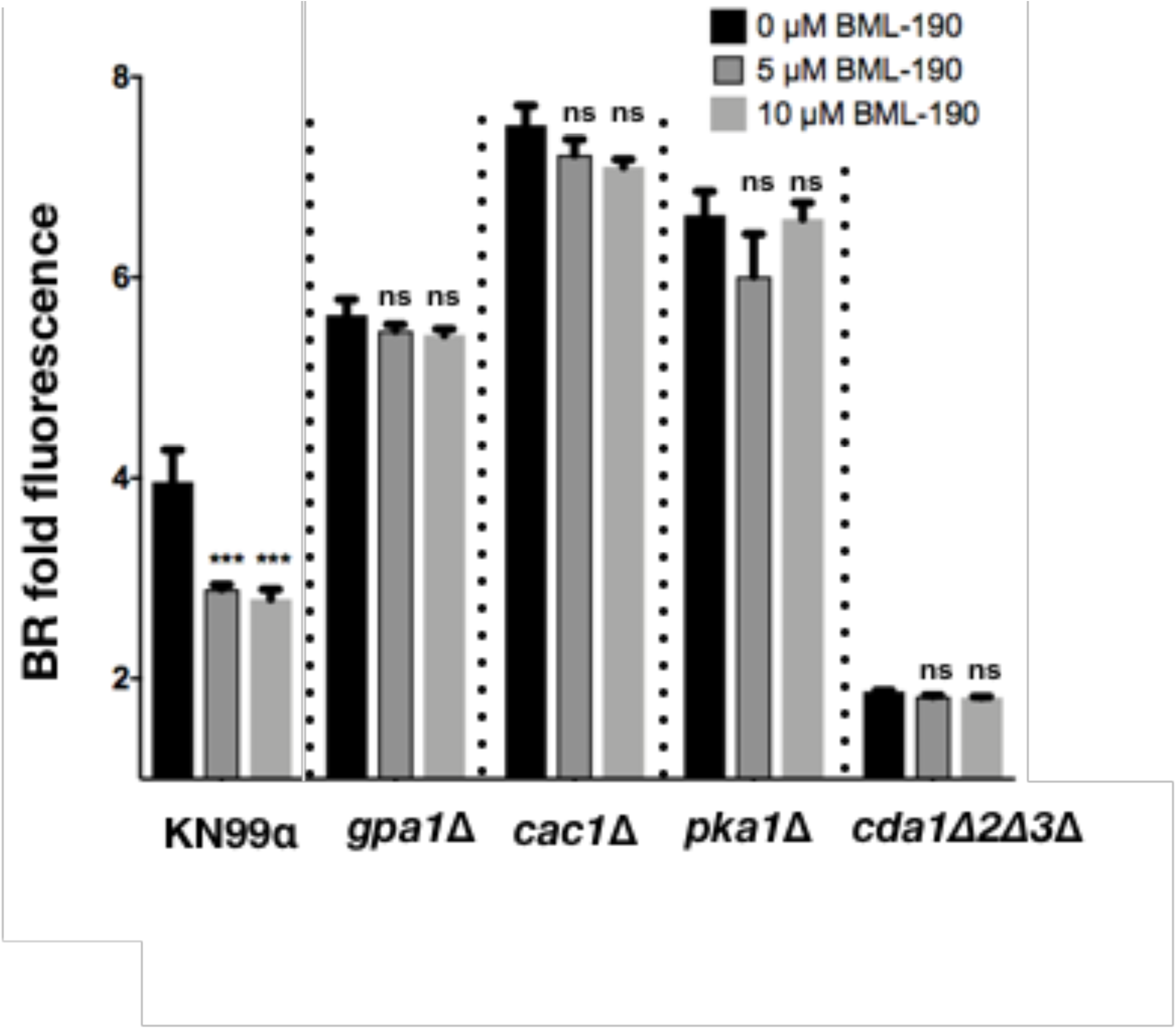
The mode-of-action for BML-190 to cause chitosan reduction is via the cAMP/PKA pathway. *C. neoformans* strains deficient in genes from the cAMP/PKA pathway were cultured with/without BML-190 for 3 days in cRPMI (0.625% HI-FBS) at 37°C and 5% CO2. These strains were stained with BR and measured for gMFI by flow cytometry. BR fold fluorescence = geometric mean fluorescence intensity (gMFI) of BR stained, BML-190 + strain/gMFI of unstained KN99α (n = 3). Means among groups were compared to strains with no BML-190 added using either a Student’s t-test or a one-way ANOVA followed by the Bonferroni multiple-comparison test, where applicable. ***, p<0.001.

**Figure 6.**
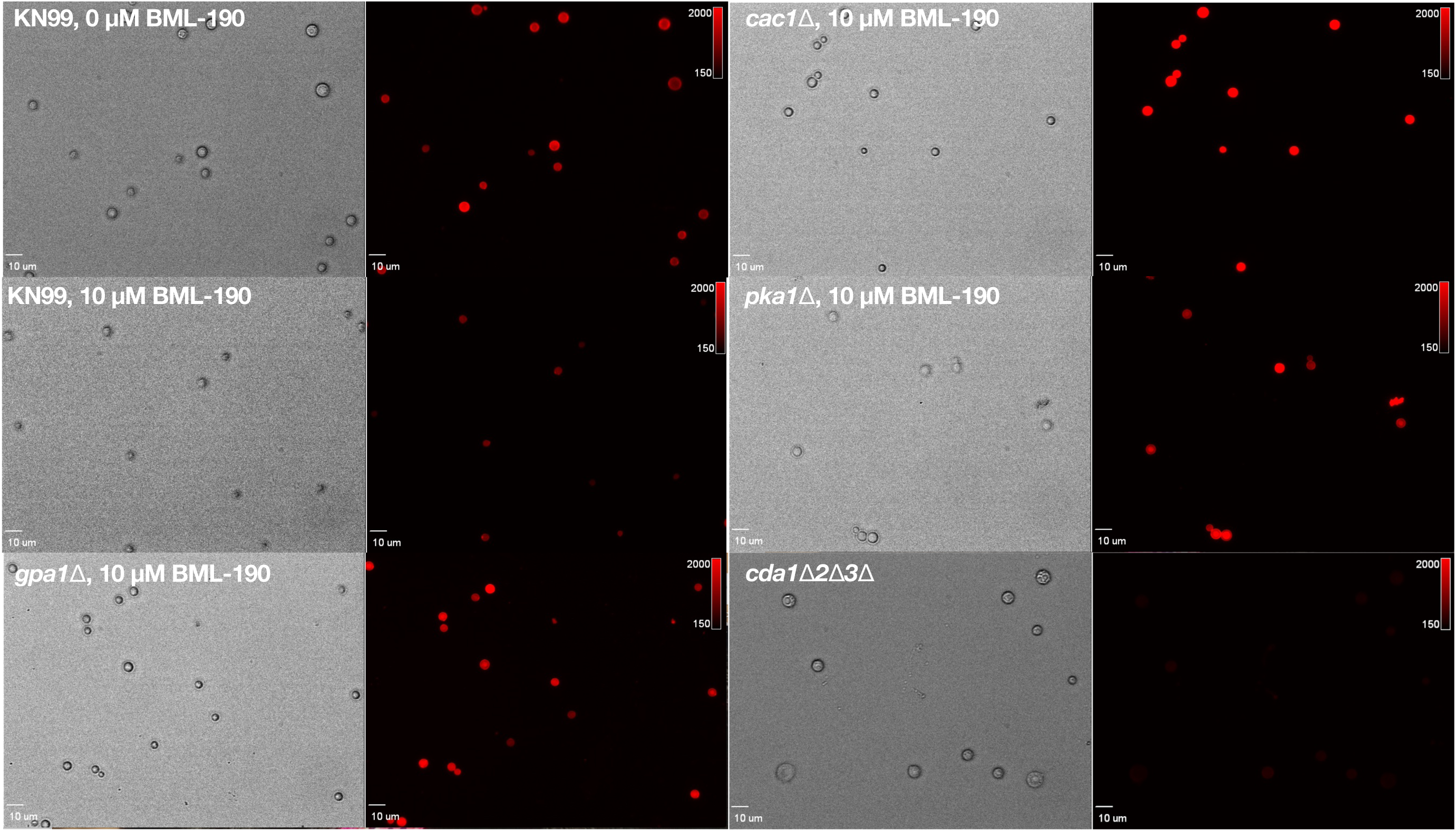
Photomicrographs showing BR fluorescence on some cAMP/PKA pathway mutants with/without BML-190. *C. neoformans* strains deficient in genes from the cAMP/PKA pathway were cultured with/without 10 µM BML-190 for 3 days in cRPMI (0.625% HI-FBS) at 37°C and 5% CO2. These strains were stained with BR and examined by epifluorescence using filter sets for TRITC (excitation 521-565 nm; emission 553-633 nm).

Pka1 is the catalytic subunit of protein kinase A, which is activated by the action of cAMP on the regulator subunit Pkr1. Similar to the upstream components of the pathway, the *pka1Δ* strain showed an increase in chitosan, and BML-190 had no effect on chitosan in this strain (Fig 5), suggesting that Pka1 may be a negative regulator of chitosan production.

Furthermore, through an examination of the other group of G-proteins in this signaling pathway, induced by BML-190, chitosan reduction is dependent on the G-protein alpha subunit (Gpa) 1, but not 2 or 3 (Fig. S5). Overall, these results support the idea that BML-190 is working through the cAMP/PKA pathway to positively and negatively regulate cAMP and chitosan production, respectively.

If cAMP is important in this pathway for regulating chitosan, we should be able to recapitulate the effects of BML-190 with the addition of cAMP to wild-type *C. neoformans*. When wild-type cells were incubated with increasing amounts of cAMP for 3 days in cRPMI, under tissue culture conditions, we observed decreasing amounts of chitosan (Fig 7). This finding is consistent with the hypothesis that BML-190 causes an intracellular accumulation of cAMP resulting in decreased chitosan. We tested this by adding increasing concentrations of cAMP to cAMP/PKA pathway mutants to see if increasing cAMP would reduce chitosan. We saw a modest, but statistically significant decrease in chitosan in wild-type and the *gpa1Δ* and *cac1Δ* at 5 mM dibutyrl-cAMP (Fig. 8). We also saw a less; albeit, significant difference in the *pka1Δ* and *gpr4Δ* strains.

**Figure 7.**
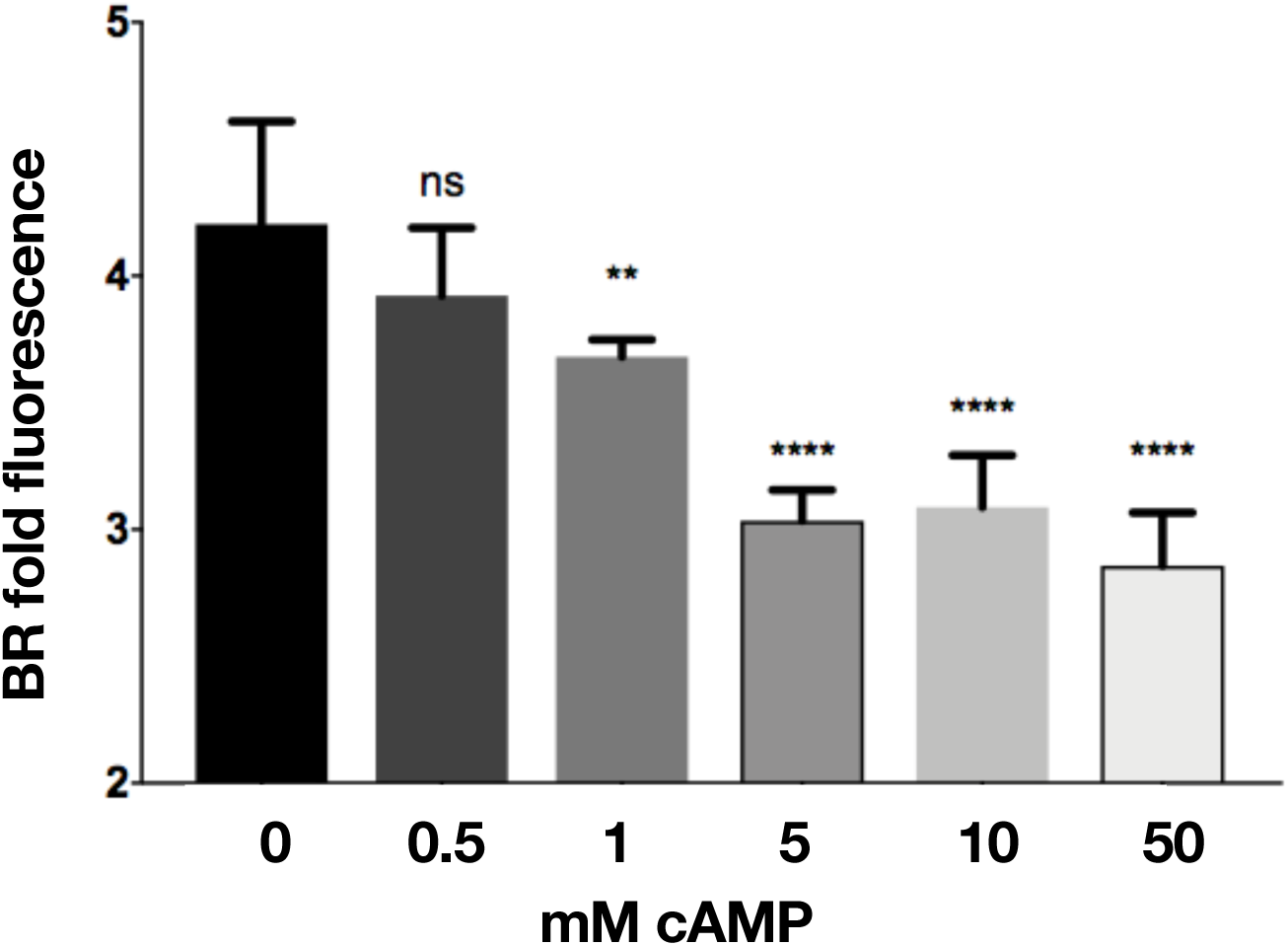
cAMP addition to *C. neoformans* results in a negative regulation of chitosan. *C. neoformans* wild-type cells were cultured with increasing concentrations of dibutyrl-cAMP for 3 days, stained with BR and measured for gMFI by flow cytometry. Means among groups (n = 5) were compared to strains with no cAMP added using a one-way ANOVA followed by the Bonferroni multiple-comparison test. **, p<0.01; ****, p<0.0001.

**Figure 8.**
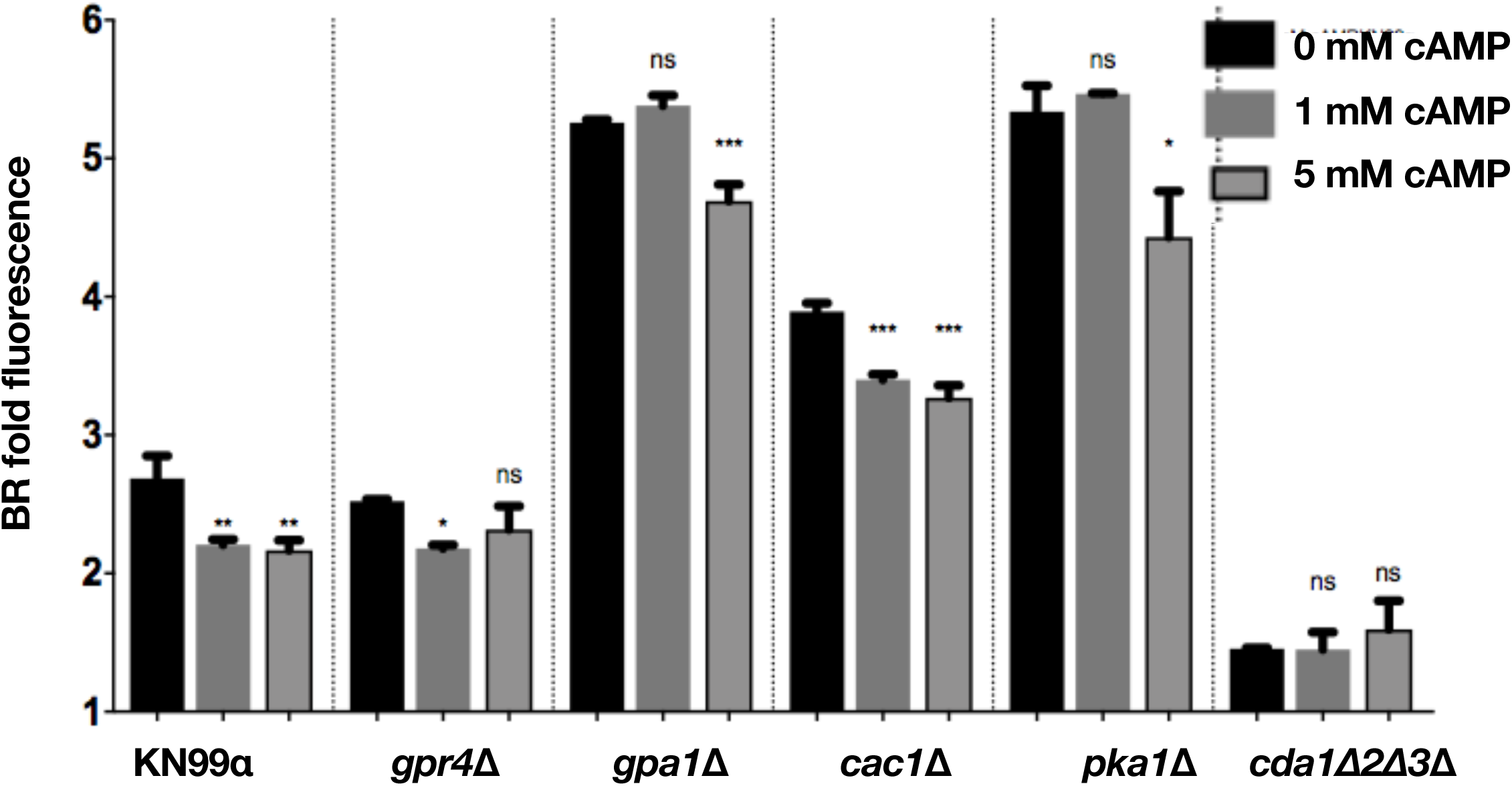
cAMP addition to *C. neoformans* cAMP/PKA pathway mutants rescues ability of pathway to reduce chitosan. A) *C. neoformans* cAMP/PKA pathway mutants were cultured with increasing concentrations of dibutyrl-cAMP for 3 days in cRPMI, stained with BR and measured for gMFI by flow cytometry. Means among groups (n = 3) were compared to strains with no cAMP added using a one-way ANOVA followed by the Bonferroni multiple-comparison test. *, p<0.05; **, p<0.01; ***, p<0.001.

The levels of BR fluorescence are higher in the *gpa1Δ* and *cac1Δ* strain, suggesting that the intracellular cAMP levels are lower in these strains compared to wild-type or the *gpr4Δ* strain. These data suggest that Cac1 and Gpa1 directly affect cAMP levels. Interestingly, the loss of Gpr4 does not appear to strongly decrease cAMP levels, based on the level of BR fluorescence. It is possible that loss of Gpr4 triggers compensating changes that allow the cAMP/Pka pathway to maintain normal levels of cAMP. Furthermore, since the *pka1Δ* strain is associated with a higher level of chitosan that is attenuated with the addition of cAMP, this implies that the regulation of chitosan is likely to be downstream of Pka1.

Finally, the role of cAMP in regulating chitosan can also be assessed by the addition of a potential agonist, which should ultimately cause a decrease cAMP that is associated with an increase in chitosan. From the primary screen of the ICCB library we discovered that there was an increase in BR fluorescence associated with the WIN 55, 212-2 compound. This compound is known to be an agonist of the cannabinoid receptors CNR1 and CNR2 contributing to a decrease in cAMP in mammalian cells (29, 35). As predicted, upon incubation of WIN 55, 212-2 compounds with wild-type *C. neoformans* we were able to show that after 3 days of incubation in cRPMI under tissue culture conditions, that there was a decrease in intracellular cAMP and an increase in chitosan as measured by BR fluorescence (Fig 9A, B). Next, we wanted to ascertain if the decrease in intracellular cAMP, caused by WIN 55, 212-2 was due to the cAMP/PKA pathway. Using previously described incubation conditions, we found that a dose-response of WIN 55, 212-2 with previously used mutant strains from the cAMP/PKA pathway resulted in an increase of chitosan in wild-type and *gpr4Δ*, a decrease of chitosan from *gpa1Δ* strains, but no effect on the *cac1Δ*, and *pka1Δ* strains (Fig 10). This suggests that WIN 55, 212-2, similar to BML-190, works through the cAMP/PKA pathway to regulate chitosan, but it does not use Gpr4 as a receptor to induce its effects. Also, interestingly, it is inducing an inverse agonist effect when Gpa1 is not expressed, suggesting that the core cAMP producing proteins can be activated by multiple signals.

**Figure 9.**
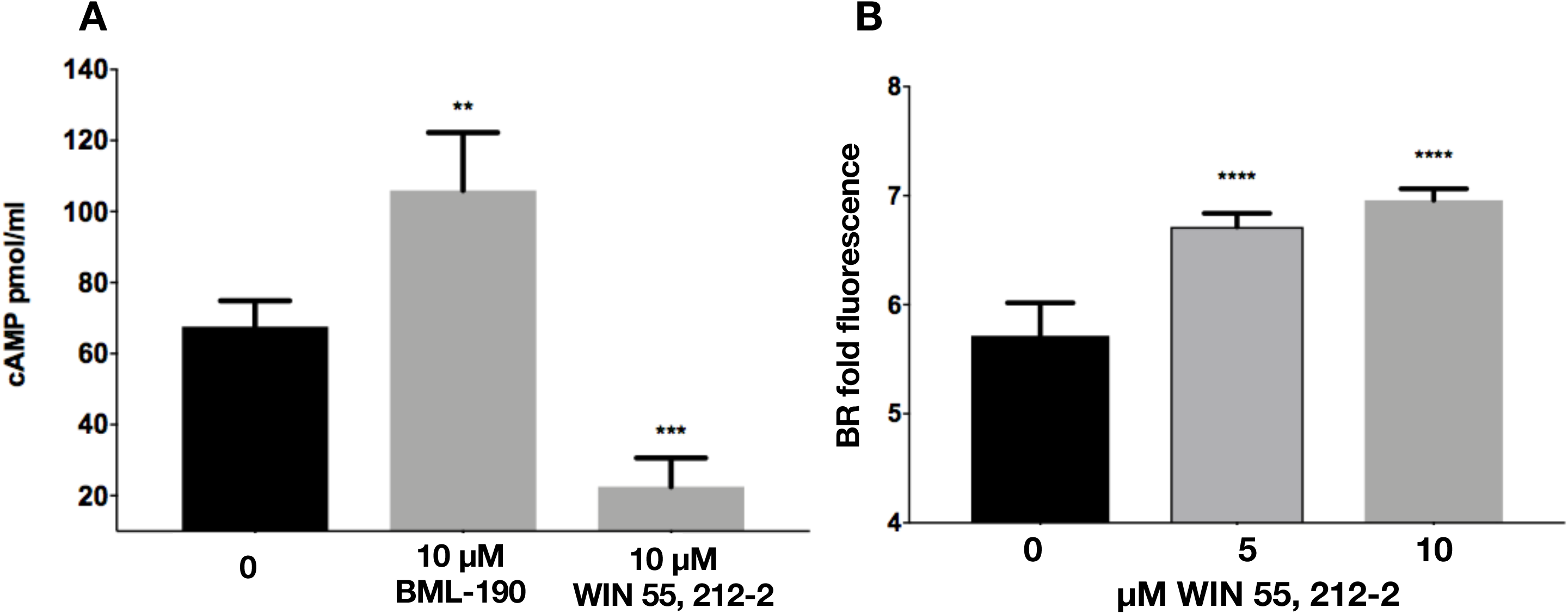
The cannabinoid receptor agonist, WIN 55, 212-2, causes a decrease in cAMP and subsequent increase in chitosan. A) Wild-type *C. neoformans* (KN99α) was cultured for 30 minutes in cRPMI with either 10 µM of BML-190 or WIN 55, 212-2 and then collected and lysed for intracellular cAMP competition ELISA (n = 5). B) A dose-response of WIN 55, 212-2-cultured *C. neoformans* (KN99α) incubated for 3 days in cRPMI, stained with BR and measured for gMFI by flow cytometry. Means among groups (n = 3) were compared to strains with no drug added using a one-way ANOVA followed by the Bonferroni multiple-comparison test. **, p<0.01; ***, p<0.001; ****, p<0.0001

**Figure 10.**
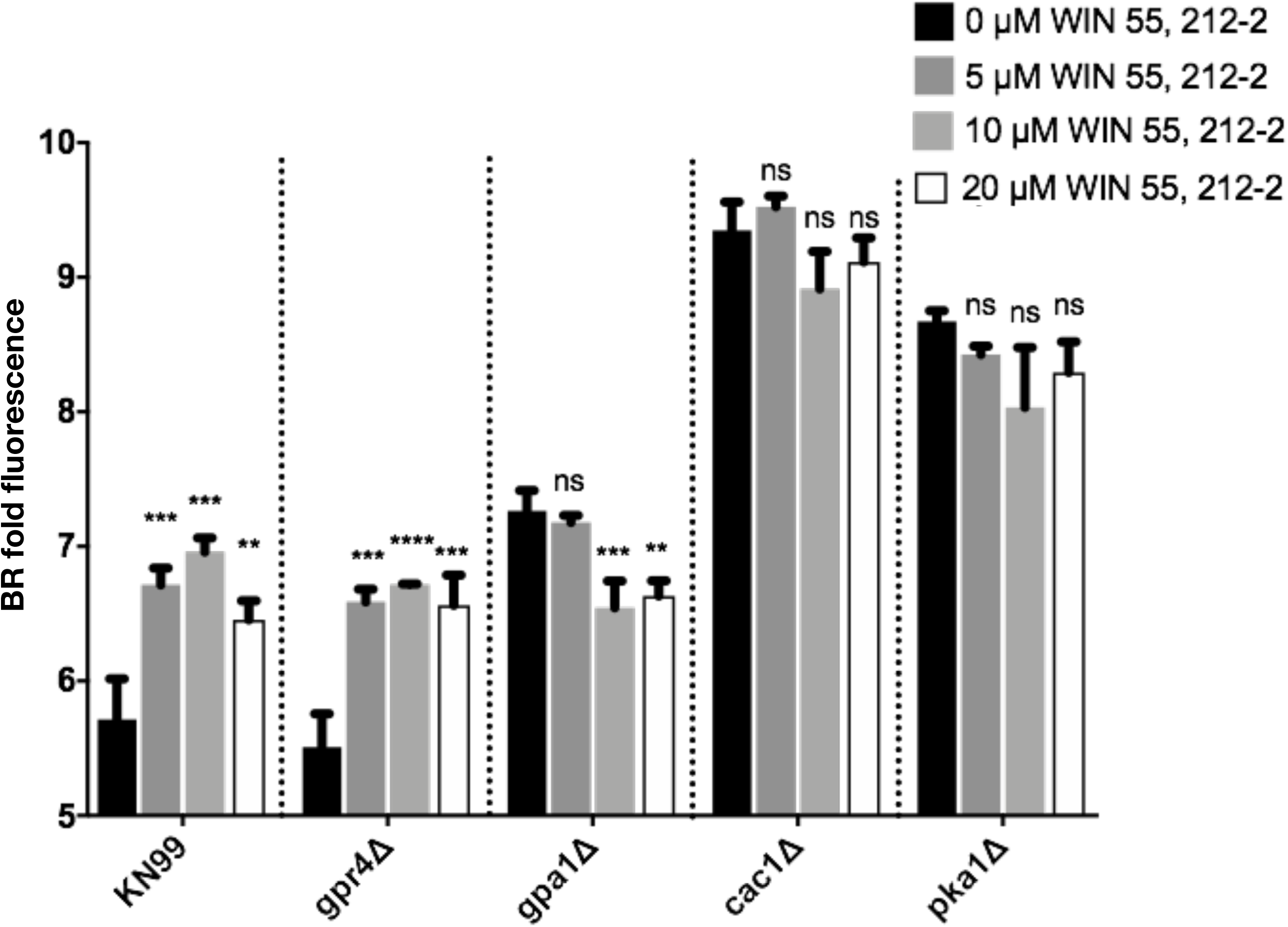
WIN 55, 212-2 regulates chitosan through the cAMP/PKA pathway. *C. neoformans* strains deficient in genes from the cAMP/PKA pathway were cultured with/without WIN 55, 212-2 for 3 days, stained with BR and measured for gMFI by flow cytometry. Means among groups (n = 3) were compared to strains with no drug added using a one-way ANOVA followed by the Bonferroni multiple-comparison test. **, p<0.01; ***, p<0.001; ****, p<0.0001

Overall, our results identify that cAMP, from the cAMP/PKA pathway is important for chitosan regulation. The pathway can be induced by BML-190, acting through the G-protein coupled receptor Gpr4 (Fig. 11). The induction by BML-190 within this pathway also requires Gpa1, Cac1, and Pka1.

**Figure 11.**
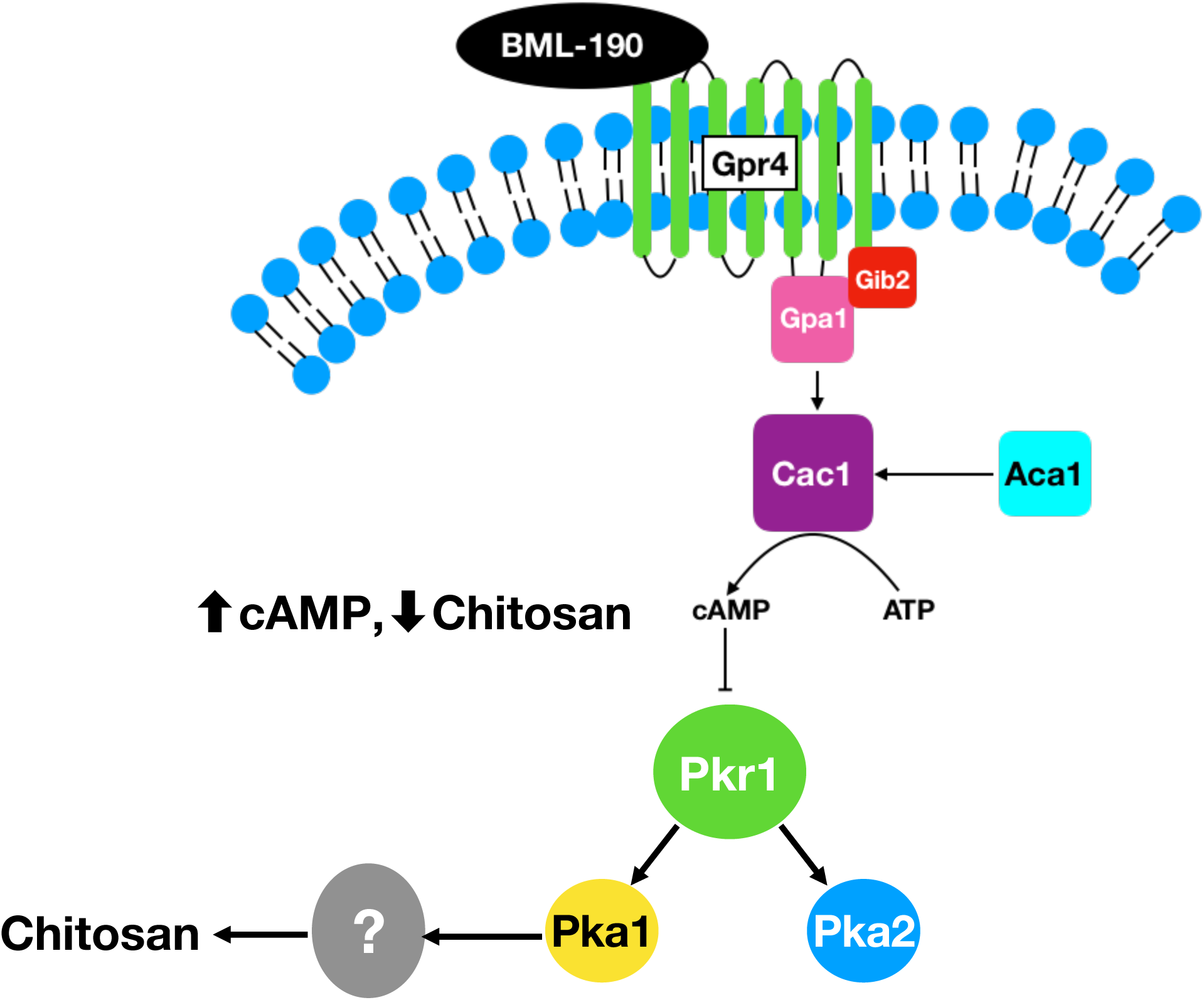
A proposed model for the target and the mode of action of BML-190 in *C. neoformans*. Pathway illustration adapted from Kronstad et. al. (33).

## Discussion

*C. neoformans* causes life-threatening meningoencephalitis in persons with AIDS. Even with antifungal therapy, mortality hovers around 30%. The current array of antifungals is insufficient to reduce worldwide fungal disease due to inadequate efficacy caused by inherent toxicities, and/or emerging drug resistance. These limitations are, in part, due to a paucity of antifungal targets. Therefore, strategies that involve the identification of novel targets that could lead to therapies with increased efficacy are needed. One approach to fulfil this goal is the targeting of factors required for growth in the host, such as chitosan production in *C. neoformans*. There is some evidence to suggest that drugs targeting virulence factors may reduce the likelihood of the development of antimicrobial resistance. The argument has been made that in the context of natural selection, if a drug does not directly kill the microbe, such as the case when targeting virulence factors, selective pressures are reduced and that the likelihood of the drug selecting for tolerant variants of the target pathogen, and promoting antimicrobial resistance in a population may also be reduced (36–38). Furthermore, because chitosan deficiency inhibits *C. neoformans* survival in the mammalian host (8, 11), even in immunocompromised hosts (data not shown), and chitosan is not produced by mammals, chitosan biosynthesis may be an ideal novel drug target for the treatment of cryptococcosis, in particular, in the context of AIDS patients who are CD4 T cell-deficient.

Through our development of a novel and robust flow cytometry primary screen for compounds that reduce chitosan production, we identified a compound, BML-190, that reduces chitosan. BML-190 is an inverse agonist targeting the G-protein coupled human cannabinoid receptors, CNR1 and CNR2. It induces adenylyl cyclase activity and causes an intracellular accumulation of cAMP (29). Since G protein signaling is known to be conserved among organisms (30), and the cAMP/PKA signaling pathway that regulates cAMP is known within *C. neoformans* (30, 34), we explored the importance of a receptor for this pathway, Gpr4, as being the target for BML-190. We treated known GPCR gene deletions, *gpr3Δ* (Fig. S5), *gpr2Δ*, *gpr4Δ* and *gpr5Δ* with BML-190 and showed that in all but the *gpr4Δ* strain BML-190 was able to cause a reduction of chitosan (Fig. 4A).

We found increased cAMP levels in wild-type cells treated with BML-190, but not in *gpr4Δ* cells (Fig. 4B). We further tested our hypothesis that cAMP levels governed by the cAMP/PKA pathway regulate chitosan levels by the use of another small molecule identified in our screen. We showed a decrease in cAMP and an increase in chitosan in cells when we treated cells with WIN 55, 212-2, an agonist towards CNR1 and CNR2 (35), supporting the involvement of the cAMP/PKA pathway (Fig. 8). Furthermore, another confirmed hit from our screen, dipyridamole is a known phosphodiesterase inhibitor that facilitates the intracellular accumulation of cAMP by preventing its hydrolysis into inactive AMP (39–40). This compound also shows a dose-dependent reduction in chitosan (Fig. S3).

*C. neoformans* protein kinase A (PKA) is a tetrameric protein that is made up of a dimeric regulatory subunit (Pkr1) and two monomeric catalytic subunits (Pka1/2). Upon cAMP targeting of Pkr1, PKA becomes activated and releases Pka1/2 which activates downstream targets by phosphorylation. It has been noted that most of the physiological effects from cAMP in fungi are mediated through targeting PKA (41). To provide intracellular cAMP for PKA targeting, Gpr4 acts as a receptor for glucose and methionine to trigger signaling through the cAMP/PKA pathway (31). When Gpr4 is physically associated with Gpa1, the adenyl cyclase enzyme (Cac1) is activated to convert ATP into cAMP (34, 42).

One important effect of PKA, through this pathway, is that it has been noted to regulate *C. neoformans* virulence. Specifically, *pka1Δ*, *gpa1Δ*, and *cac1Δ* mutant strains, have reduced virulence along with reduced melanin and capsule volume (34). Furthermore, PKA is known to promote cell proliferation (43) and negatively regulates cAMP, acting as a negative feedback loop for PKA activity via its activation of phosphodiesterase (Pde1; 40). Since we showed that chitosan production is inversely related to cAMP levels (Fig 2C&D, Fig 8) and that there is no decrease in chitosan in the mutant strains of *pka1Δ*, *gpa1Δ*, and *cac1Δ* with increasing concentrations of BML-190 (Fig 5), it can be further supported that the mode of action of BML-190 is the cAMP/PKA pathway. Interestingly, the chitosan levels are higher in these mutant strains compared to wild-type with/without BML-190. Since we have noted that there is an inverse relationship between chitosan and cAMP production, lower cAMP should produce greater chitosan. This would be the case in *gpa1Δ*, and *cac1Δ* mutants since these are proteins noted to be important for cAMP formation. However, this does not explain the higher chitosan production in *pka1Δ* mutants which should have relatively high amounts of cAMP (40) because Pde1 is not activated by Pka1 to degrade cAMP. One possibility is that cAMP is negatively regulated by additional mechanisms in addition to Pde1. Also, through our dose-response of adding exogenous cAMP to *pka1Δ*, in turn, causing a loss of chitosan (Fig 8), it suggests that cAMP levels are indeed low in the examined cAMP/PKA deletion strains which conforms to our hypothesis of the inverse relationship between cAMP and chitosan. It’s possible that *pka1Δ* mutants grown under optimal growth conditions (i.e., YPD growth conditions) show greater cAMP levels (40) than those grown under host conditions (i.e., cRPMI growth at 37°C and 5% CO_2_) as was done for this study.

As previously described, PKA is a well-described mediator of the effects of cAMP on inducing virulence factors in *C. neoformans*. This has been identified to include their ability to proliferate under host conditions (37°C; 40), produce an antiphagocytic polysaccharide capsule, and antioxidant melanin (34, 41–42). Surprisingly, BML-190 not only reduces chitosan, but our data suggests it also reduces capsule volume and melanin (Fig. S6; Fig. S7). Based upon our evidence that BML-190s mode of action is the cAMP/PKA pathway, with cAMP targeting PKA to induce a mechanism downstream to cause a reduction in chitosan, it is also possible that this downstream mechanism is also responsible for causing a reduction in the other virulence factors of capsule volume, melanin, as well as decreased growth at 37°C (Fig 2). Another possibility is that crosstalk is occurring between another pathway involved in regulating virulence factors such as the cell wall integrity (CWI) pathway, which we have shown interacts with the cAMP/PKA pathway (44). However, this mechanism has yet to be elucidated.

In conclusion, the paucity of drug targets and the lower efficacy of current treatments for cryptococcosis, a prominent cause of morbidity and mortality in AIDS patients, additional treatments must be discovered. To help increase efficacy, drugs and their targets must be found that are targeting the pathogen and are non-toxic to the host. By targeting chitosan of the pathogen, we fulfil that criteria. However, because BML-190 also has been shown to have effects on cannabinoid receptors, it is unlikely to be developed further as an anti-cryptococcus drug, but additional screening for reduced chitosan using the flow-cytometry cell-based screen could identify compounds that do not impact the mammalian host. Furthermore, by the drug disarming the pathogen, as opposed to killing it within the host, it is possible that there may be a reduced effect of natural selection by the drug on the pathogen, in turn, a potential reduction in their antimicrobial resistance to the drug, but this remains to be determined.

Our discovery that BML-190 negatively regulates chitosan via the cAMP/PKA signaling pathway also suggests that our screening strategy can identify more compounds from additional drug libraries that will regulate chitosan production from *C. neoformans* to further identify key genes and proteins critical for the production and regulation of chitosan and cell wall biosynthesis.

## Methods and Materials

### Cryptococcus strains and growth conditions

*C. neoformans* strains KN99*α*, the mCherry reporter strain (KN99mCH; 14), and those engineered that differentially express chitosan, *cda1Δ*, *cda2Δcda3Δ*, and *cda1Δcda2Δcda3Δ* (referred to as *cda2Δ3Δ* and *cda1Δ2Δ3Δ*, respectively, throughout the manuscript), have been described previously (9). Additional strains used included those derived from KN99*α* with single deletions in G-protein coupled receptors, G-protein alpha subunits, and cAMP/PKA pathway that were part of the deletion library generated by Hiten Madhani at UCSF and obtained from the Fungal Genetic Stock Center (FGSC; 15). These deletion strains (followed by gene ID) were: *gpr2*Δ (CNAG_01855), *gpr3Δ* (CNAG_03846), *gpr4Δ* (CNAG_04730), *gpr5Δ* (CNAG_05586), *gpa1Δ* (CNAG_04505)*, gpa2Δ* (CNAG_00179)*, gpa3Δ* (CNAG_02090)*, cac1Δ* (CNAG_03202), and *pka1Δ* (CNAG_00396). All strains were streaked onto yeast-peptone-dextrose (YPD) agar plates from −80°C stocks. After the development of colonies, strains were further propagated by their addition to YPD broth and grown for 72 hours at 30°C and 300 RPM.

### Flow cytometry cell-based phenotypic screening assay

We screened 480 compounds from the Institute of Chemistry and Cell Biology (ICCB) Known Bioactives library (Enzo Life Sciences) using our optimized cell-based flow cytometry phenotypic assay. We used a Biomek FX workstation to add 1µl of each compound to columns 2-11 of 96-well sterile round-bottom tissue culture plates in duplicate. For the controls, 1µl DMSO was added to Columns 1 and 12. The screening strain (*cda2Δ3Δ*) and the chitosan-deficient strain (*cda1Δ2Δ3Δ*) were initially cultured in the complete growth medium YPD, washed 2x in 1x PBS, counted by hemocytometer, and resuspended at 5×10^5^ cells/mL in complete RPMI (cRPMI; 0.625% HI-FBS). We added 100 µl of cells per well, for a final DMSO concentration of 1%, which ensured the solubility of the ICCB compounds. We found minimal impact on growth of either strains under these conditions (data not shown). Plates were incubated for 3 days at 37°C with 5% CO_2_. After incubation, we washed the cells with 1X PBS and stained them with Cibacron Brilliant Red 3B-A (BR) for 20 minutes at a final concentration of 5 µg/ml in PBS, pH 2. We resuspended one well of the screening strain in 1X PBS at a pH = 2 without BR as an unstained control to measure background fluorescence. Cells were then pelleted via centrifugation at 1000 *x g* for 10 minutes at room temperature in the plates, flick decanted, and fixed by resuspension in 1% paraformaldehyde (PFA) followed by overnight incubation at 4°C. Prior to flow cytometry we centrifuged the plates, flick-decanted and resuspended the cells in flow cytometry buffer (FCB buffer; 0.5% BSA+2mM EDTA+1x PBS). All plates were then analyzed for multiple parameters using a BD Biosciences LSRFortessa X-20 flow cytometer equipped with a High-throughput Sampler (HTS), which enables automated sample acquisition from 96- and 384-well microtiter plates. BR fluorescence was detected by excitation with a Yellow-Green laser (561 nm) and emission collection with the PE channel (586/15nm). We were able to complete the analysis of twelve plates (1152 wells = compounds + controls) in 6.5 hours. The average Z-prime for these 12 plates was 0.77, indicative of a robust screening assay (16).

### 3-methyl-2-benzothiazolone hydrazone hydro-chloride (MBTH) chitosan assay

*C. neoformans* strains were grown in 25 mL of cRPMI (0.625% HI-FBS) at 5×10^5^ cells/mL under tissue culture conditions for 72 hours at 37°C and 5% CO_2_. Cells were then pelleted into preweighed 15 mL conical centrifuge tubes by centrifugation (3000 × *g* for 10 minutes). Supernatant was aspirated and then samples were resuspended with 1x PBS and then cells were again pelleted. Pelleted cells were then weighed after being freeze-dried to normalize chitosan measurements to dry weights. To reduce background chromogens, cells were alkaline extracted by the addition of 10 mL of 6% KOH to cell pellets followed by incubation at 80°C for 90 minutes and periodic mixing. Pellets were then washed twice with 10 mL 1x PBS followed by two washes in ultrapure water. Samples were then all normalized to their initial dry weights by addition of ultrapure water to make all samples 10 mg/mL.

For MBTH colorimetric determination of chitosan, samples of 100 µl (1 mg) were diluted 1:1 with 1 M HCL. Then, 400 µl of 0.36 M of sodium nitrite was added, vortexed, and incubated for 15 minutes at room temperature. Next, 200 µl of 1.1 M of ammonium sulfamate was added, vortexed, and incubated for 5 minutes at room temperature. This was then followed by the addition of 200 µL of 0.011 M MBTH, followed by vortexing, and incubation for 30 minutes at 37°C. To induce blue color formation, 200 µL of 0.018 M of iron(III) chloride hexahydrate was added and vortexed, followed by incubation for 5 minutes at 37°C. Samples were cooled to room temperature, centrifuged at 14,000 RPM for 2 minutes. 100 µL of supernatant from each sample to wells of a flat-bottom well microtiter plate and absorbance measured at 650 nm. A standard curve was prepared from 2-fold serial dilutions of 2 mM GlcN for the assessment of chitosan concentration of samples.

### cAMP measurements

We grew *C. neoformans* strains overnight in YPD, washed the cells 2x in 1X PBS and added them at a concentration of 5 × 10^8^ cells/ml to T-175 tissue culture flasks containing 10 ml of cRPMI (0.625% HI-FBS) with final concentration of 1% DMSO with/without testing compound. We incubated the flasks for 30 minutes at 37°C and 5% CO_2_, then collected media from each flask and centrifuged at 3000 *x g* for 10 minutes at 4°C. We aspirated the media and resuspended the cell pellets in 2 ml of ice cold 0.1 M HCl, transferred 1.5 ml to 2 ml screw top microtubes (Sarstedt Inc., Numbrecht, Germany) on ice that contained 0.5 mm zirconia/silica beads (Biospec. Products, Bartesville, OK, USA). Cells were homogenized using a Mini-Beadbeater-16 (Biospec. Products) at 75% of maximum agitation at 4°C for 3 rounds of 2 minutes on and 2 minutes off. We transferred the homogenized samples to new tubes and stored at −20°C. We thawed the samples on ice, centrifuged at 3000 *x g* for 10 minutes and transferred the supernatants to fresh tubes. We measured cAMP concentrations using a cAMP competitive ELISA kit according to the manufacturer’s instructions (Sigma-Aldrich, St. Louis, MO, USA). Samples were added to ELISA wells as 100 µl aliquots (2.5 × 10^8^ cell lysate equivalent) and were not acetylated prior to measurements.

### BR Staining of *C. neoformans* for fluorescent microscopy

We dissolved 10 mg/mL of BR in ultrapure water. Cryptococcus cells grown in YPD were washed once in PBS (1000 *x g* for 10 minutes) and diluted to 1 × 10^7^ cells/mL PBS. Cells were then pelleted and resuspended in 5 µg/mL of BR in PBS (pH = 2) followed by incubation at room temperature for 20 minutes in the dark. We washed the cells with 1X PBS and resuspend at 4×10^7^ cells/ mL in PBS. We transferred 10 uL to a microscope slide, covered with a cover slip and sealed with nail polish. Slides were then examined by epifluorescence using filter sets for TRITC (excitation 521-565 nm; emission 553-633 nm).

### Capsule measurements

Wild-type *C. neoformans* (KN99α) cells were grown in cRPMI (0.625% HI-FBS) and 0.1% DMSO with/without 20 µM BML-190 for 3 days at 37°C and 5% CO_2_. Cells were then washed in 1xPBS and resuspended in 1/4 dilution of India ink (50 µl) and then added (10 µl) to a microscope slide. The diameters of whole cell and cell body were measured from 60 different cells within each biological replicate. Volume of a sphere was determined from those diameters. Capsule volume was determined by subtracting the volume of the cell body from the volume of the whole cell. Means between groups (n = 3) were compared using a Student’s t Test.**, p< 0.01.

### Melanin measurements

Wild-type *C. neoformans* (KN99α) was grown in cRPMI (0.625% HI-FBS) and 0.1% DMSO with/without 20 µM BML-190 for 3 days at 37°C and 5% CO_2_ (n = 3). Cells were then washed in 1xPBS. Then, 1 × 10^8^ cells (5 × 10^7^/ml), from each condition above, were added to 2 ml of glucose-free asparagine medium (1 g/liter L-asparagine, 0.5 g/liter MgSO4 7H2O, 3 g/liter KH2PO4, and 1 mg/liter thiamine, plus 1 mM L-3,4-dihydroxyphenylalanine (L-DOPA)) for 7 days at 300 RPM and 30°C. Samples were then spun down at >600 × *g* for 10 minutes. Pellets were then transferred to 96-well flat-bottom microtiter plate and then photographed.

### Statistical analysis

All data from experiments are from greater than or equal to three independent experiments. Error bars represented standard deviation. Inferential statistics used at least one of the following relevant parametric analyses: A two-tailed unpaired Student’s t test, a one-way ANOVA with Bonferroni multiple comparison test, and/or a Pearson’s r correlation analysis. For all experiments, *\*p* ≤ 0.05, \**\*p* ≤ 0.01, *\*\*\*p* ≤ 0.001, and \*\*\**\*p* ≤ 0.0001 indicated statistically significant differences. All statistical analysis was completed on Prism 7 for Mac OS X software (Version 7.0c).

## Data Sharing

The screening data will be deposited in the publicly available database https://pubchem.ncbi.nlm.nih.gov prior to publication.

## Acknowledgements

We thank Michael Prinsen for his technical assistance and Rajendra Upadhya, Camaron Hole, and Abigail Ragsdale for their helpful discussions about the paper.

## Supplementary Figures

**Figure S1.**
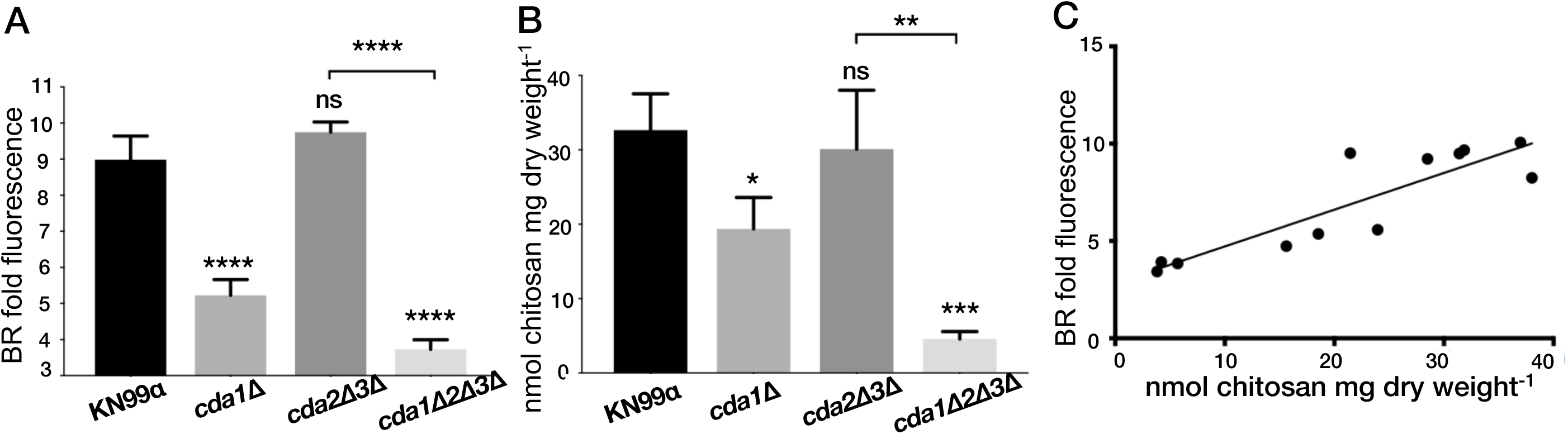
Known differential expression of chitosan by *C. neoformans* strains was identified by flow cytometry. A) Flow cytometry generated BR fold fluorescence shows a correlation with the B) MBTH Chitosan assay when identical strains of *C. neoformans* are used for both assays. Means among groups were compared to KN99α group using a one-way ANOVA followed by the Bonferroni multiple-comparison test. *, p<0.05; **, p< 0.01; ***, p<0.001, and ***, p<0.0001. C) Based upon a Pearson’s r correlation analysis, chitosan measured by flow cytometry and the MBTH strongly and statistically covary with one another (r = 0.8789). N = 12, critical r = 0.576, p < 0.001. Both conditions are grown in complete RPMI for 3 days at 37°C and 5% CO2. BR fold fluorescence = geometric mean fluorescence intensity (gMFI) of BR stained strain/gMFI of unstained KN99α strain.

**Figure S2.**
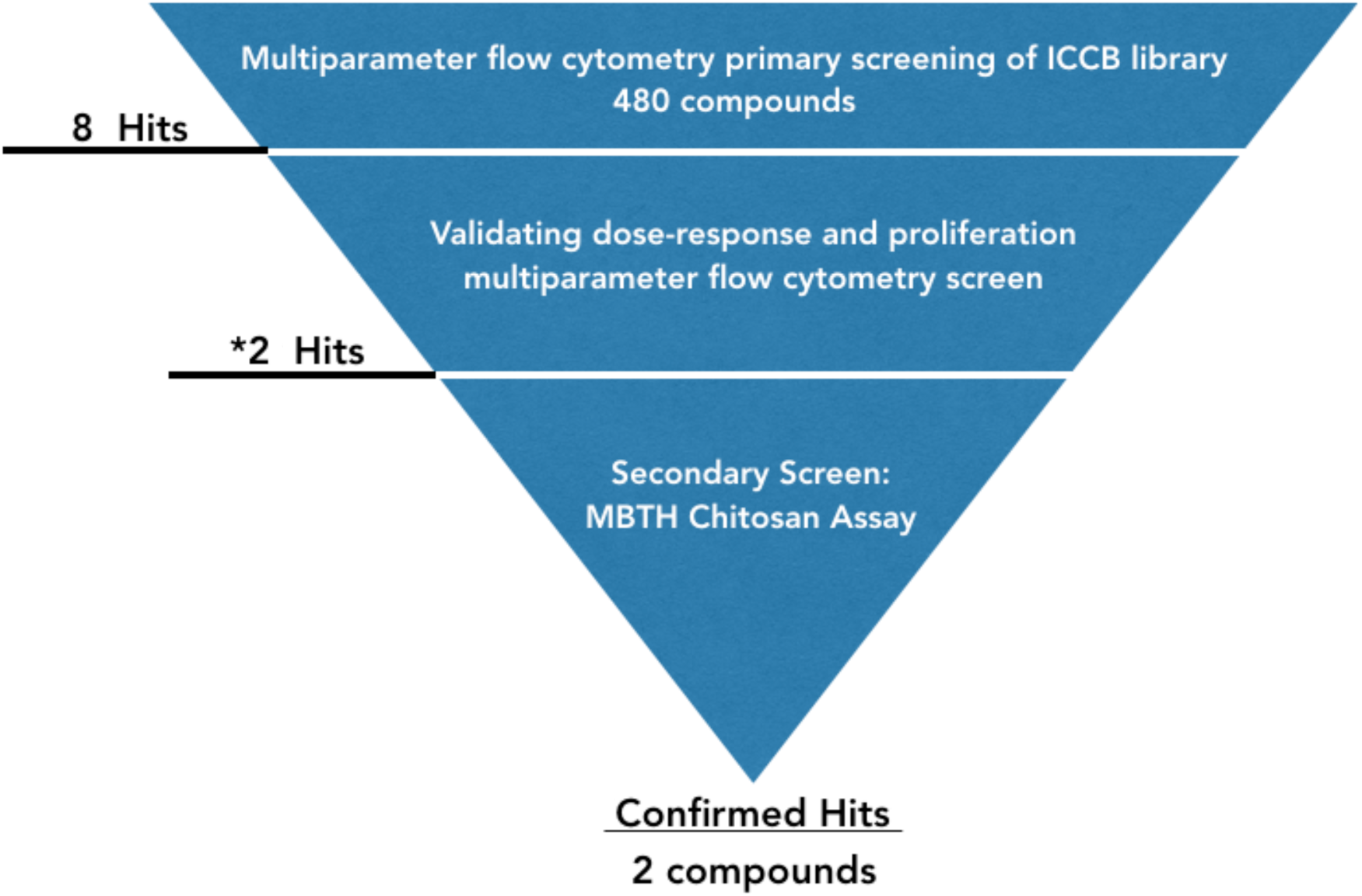
Screening strategy funnel for inhibitors of chitosan production from *C. neoformans*. An overview of the the primary cell-based phenotypic screen of the 480 compounds from the ICCB library. From our primary screen, we identified 8 hits. Of those 8 hits, 6 were further validated to identify 2 hits. Those hits were identified as confirmed hits by our MBTH Chitosan assay: dipyridamole and BML-190.

**Figure S3.**
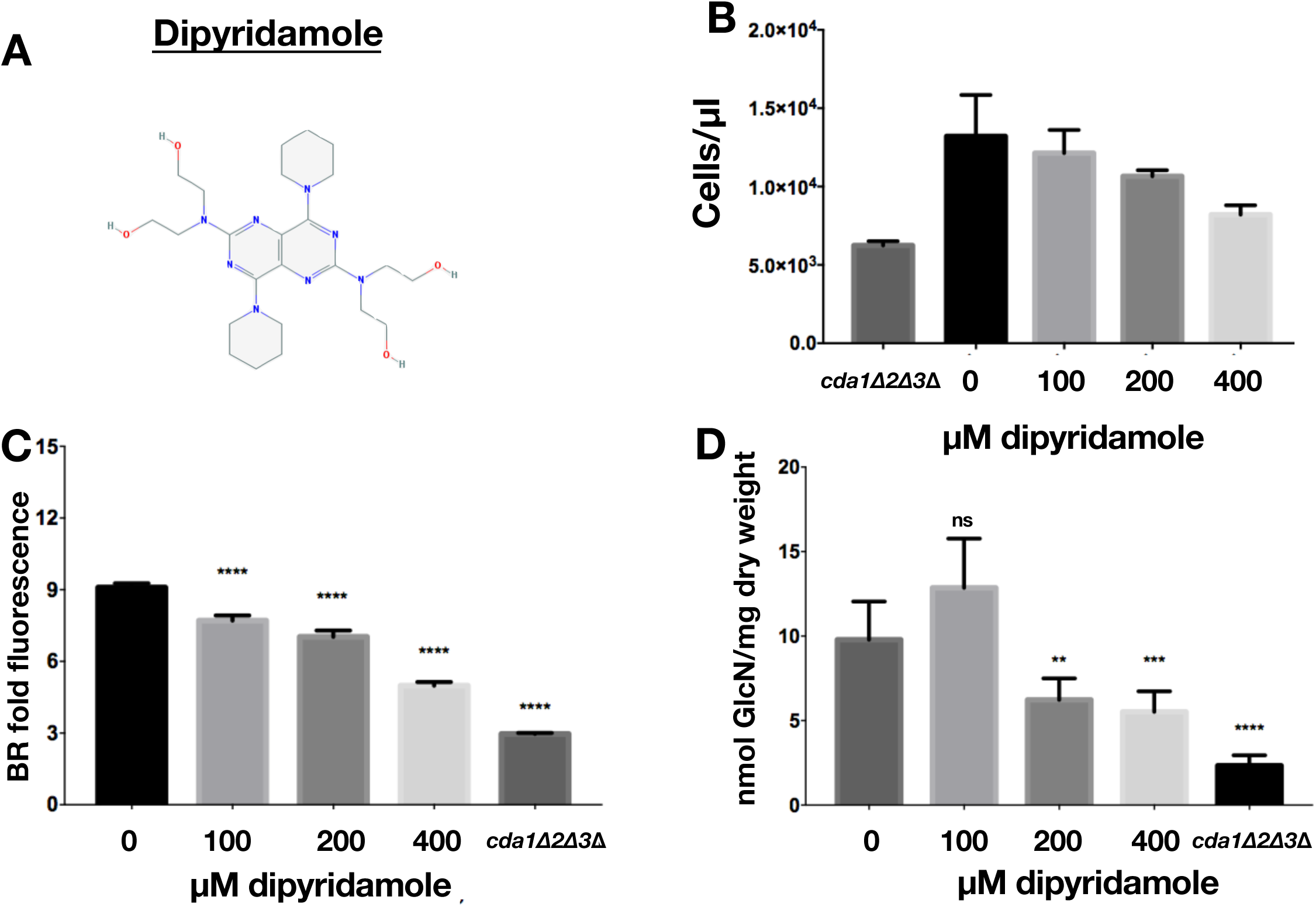
The phosphodiesterase inhibitor, dipyridamole reduces chitosan in *C. neoformans*. A) Chemical structure of dipyridamole. B) *C. neoformans* concentrations determined by flow cytometry using counting beads with dipyridamole concentrations between 100 µM to 400 µM that were incubated for 3 days in cRPMI. C) A dose-response of dipyridamole-cultured *C. neoformans* (KN99α) incubated for 3 days in cRPMI, stained with BR (BR) and measured for gMFI by flow cytometry. C) A dose-response of dipyridamole-cultured *C. neoformans* (KN99α) incubated for 3 days in cRPMI and chitosan content determined using the MBTH assay. Means among groups (n = 3) were compared to cda1Δ2Δ3Δ (B) or KN99α (C-D) with no BML-190 added using a one-way ANOVA followed by the Bonferroni multiple-comparison test. *, p<0.05; **, p<0.01; and ****, p<0.0001.

**Figure S4.**
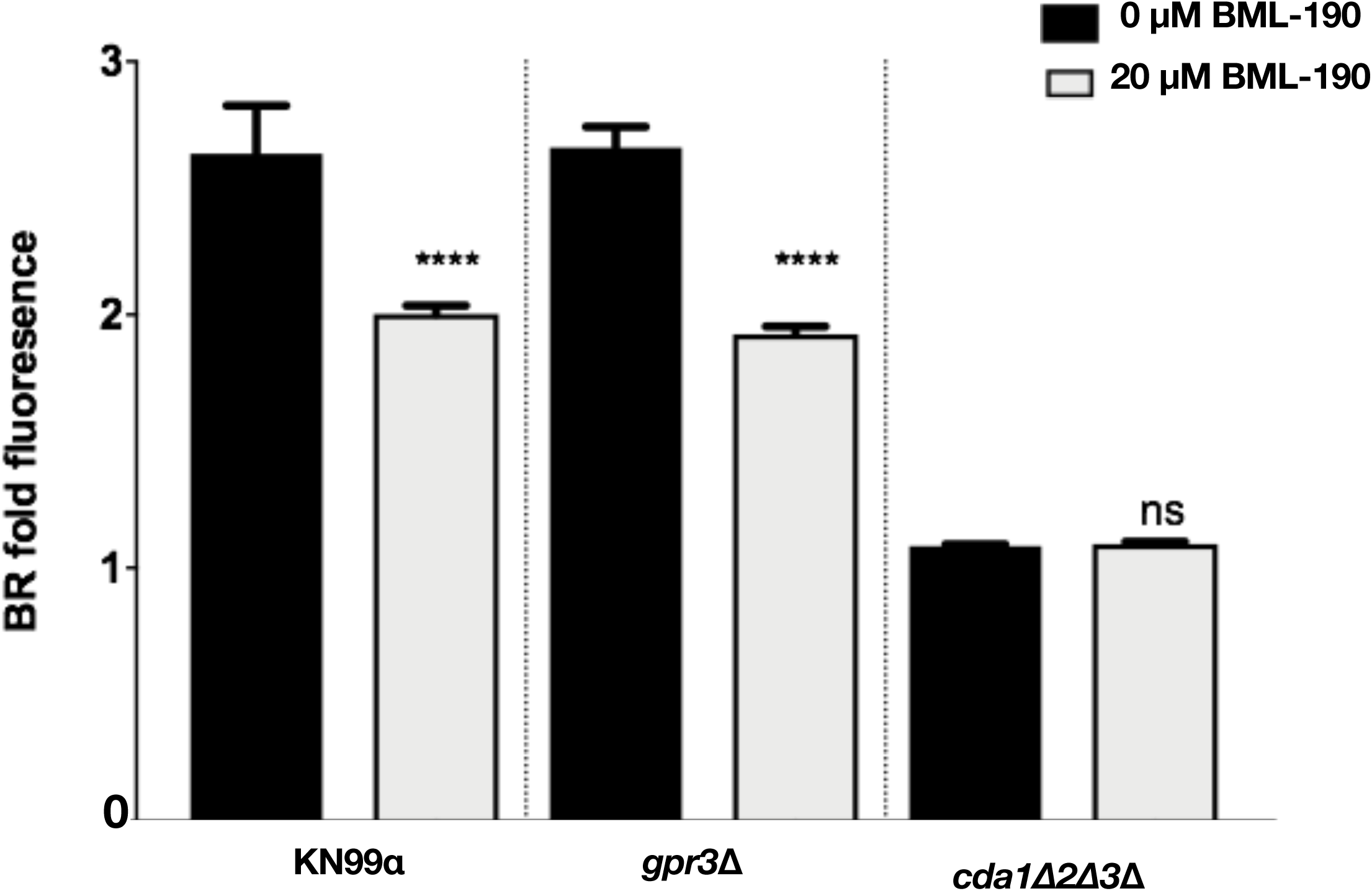
Gpr3 is not a receptor for BML-190. *C. neoformans gpr3Δ* were cultured with/without BML-190 (20 µM) for 3 days in cRPMI, stained with BR and measured for gMFI by flow cytometry. Means among groups (n = 6) were compared to strains with no BML-190 added using a Student’s t Test, ****, p<0.0001.

**Figure S5.**
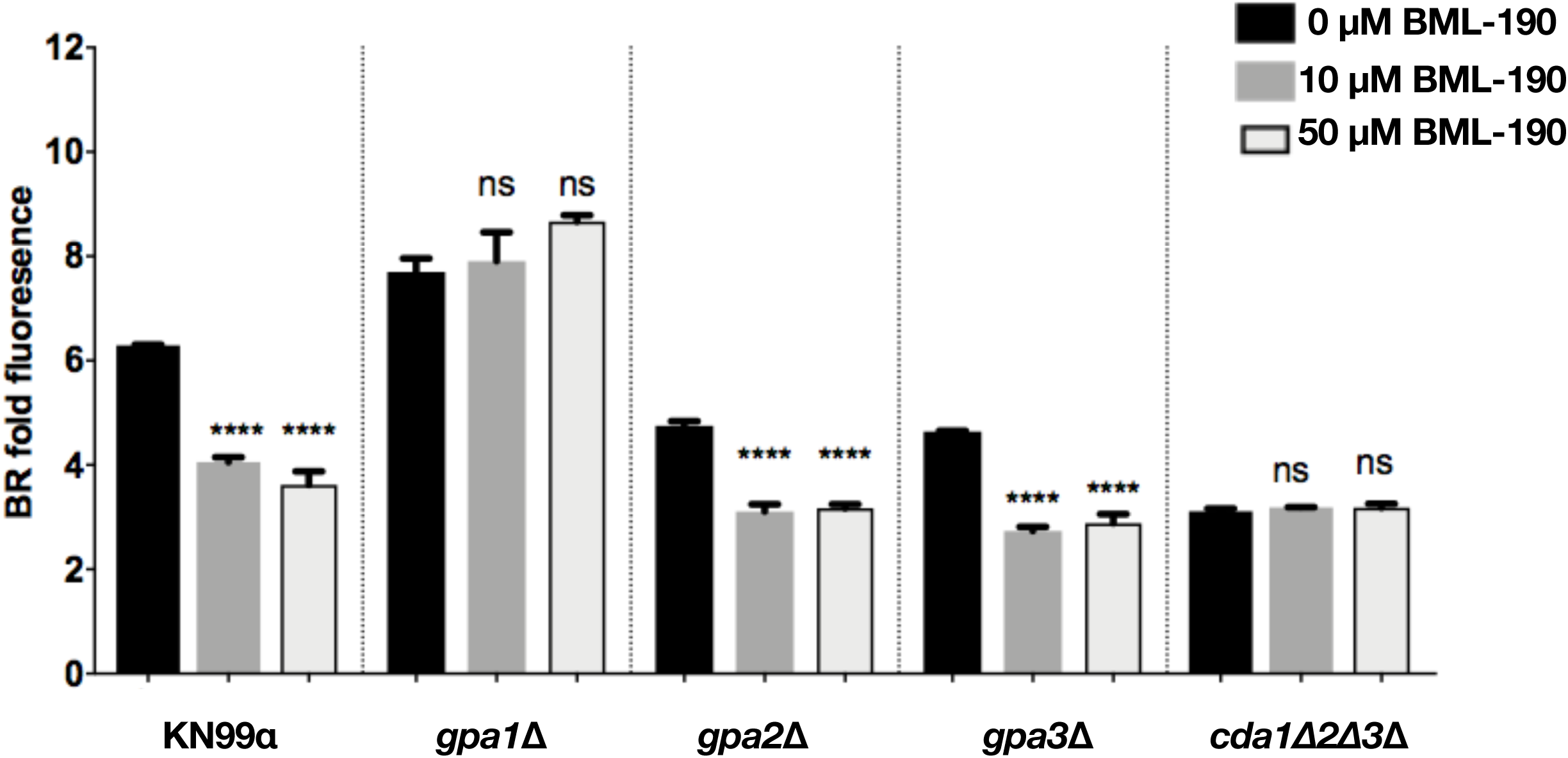
BML-190 causes chitosan reduction by working through the G-protein alpha subunit 2 Gpa1. C. neoformans gpa-deficient strains were cultured with/without BML-190 (10 µM and 50 µM) for 3 days in cRPMI, stained with BR and measured for gMFI by flow cytometry. Means among groups (n =4) were compared to strains with no BML-190 added using a one-way ANOVA followed by the Bonferroni multiple-comparison test. ****, p<0.0001.

**Figure S6.**
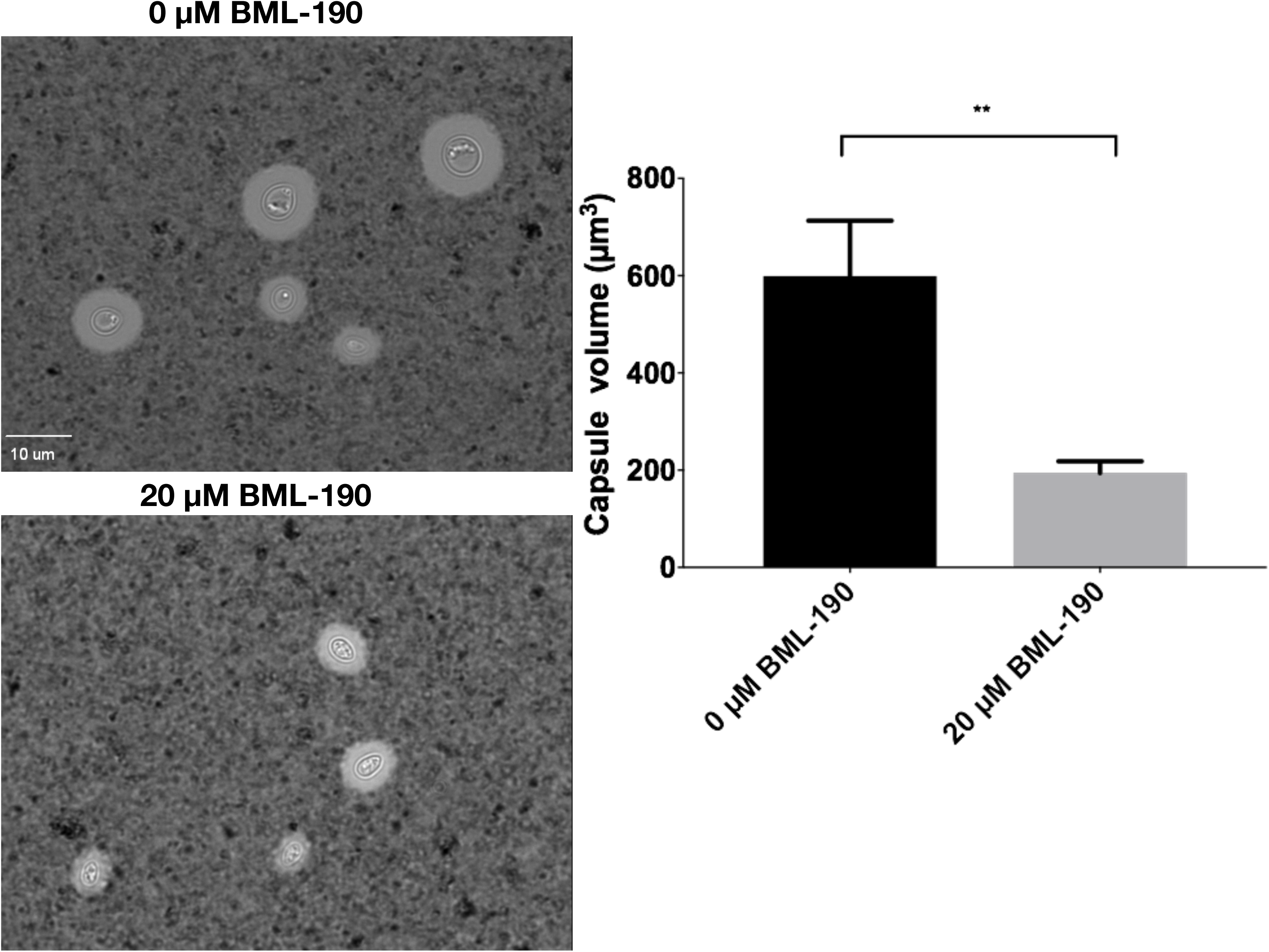
BML-190 causes a decrease in capsule volume. Wild-type *C. neoformans* (KN99α) was grown in cRPMI (0.625% HI-FBS) and 0.1% DMSO with/without 20 µM BML-190 for 3 days at 37°C and 5% CO_2_. Cells were then washed in 1xPBS and resuspended in 1/4 dilution of India ink (50 µl) and then added (10 µl) to microscope slide. The diameters of whole cell and cell body were measured from 60 different cells within each biological replicate. Volume of a sphere was determined from those diameters. Capsule volume was determined by subtracting the volume of the cell body from the volume of the whole cell. Means between groups (n = 3) were compared using a Student’s t Test.**, p< 0.01.

**Figure S7.**
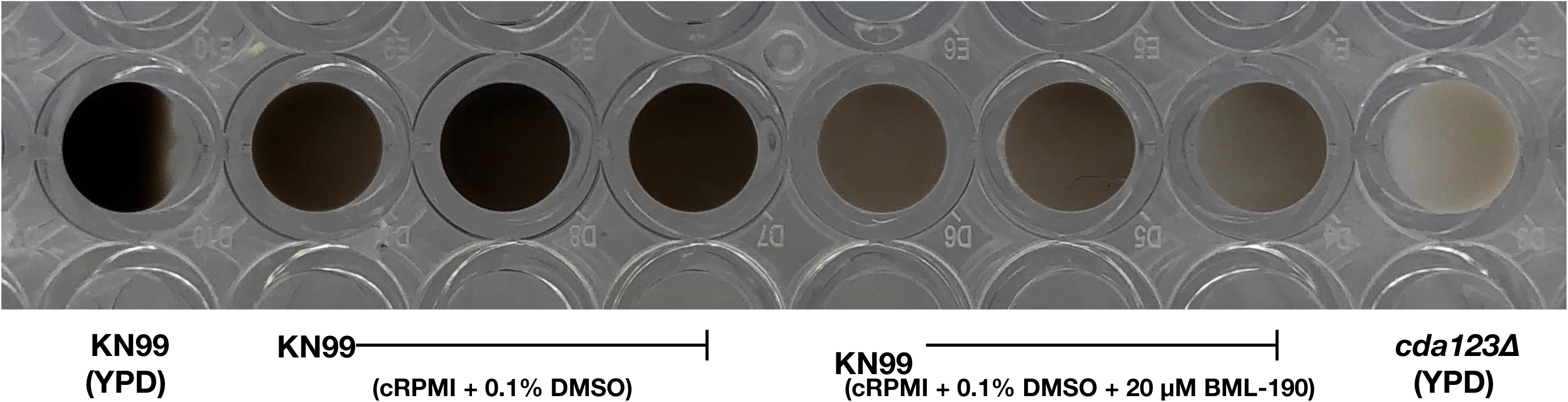
BML-190 causes a decrease in *C. neoformans* melanin. Wild-type *C. neoformans* (KN99α) was grown in cRPMI (0.625% HI-FBS) and 0.1% DMSO with/without 20 µM BML-190 for 3 days at 37°C and 5% CO_2_ (n = 3). Cells were then washed in 1xPBS. 1 × 10^8^ cells, from each condition above, were added to 2 ml of glucose-free asparagine medium for 7 days at 30°C.

